# Mechanical and Growth Anisotropy in *Chara corallina*: Challenging Green’s Hypothesis

**DOI:** 10.64898/2026.02.09.704413

**Authors:** Weiyuan Kong, Antonio Mosciatti, Jerome Boulanger, Guillaume Marrelec, Thierry Savy, Etienne Couturier

**Affiliations:** Laboratoire Matière et Systèmes Complexes, Université Paris Diderot CNRS UMR 7057, 10 rue Alice Domont et Léonie Ducquet, 75205 Paris Cedex 13, France; MRC Laboratory of Molecular Biology, Francis Crick Avenue, Cambridge, CB2 OQH, UK; Laboratoire d’imagerie biomédicale, LIB, Sorbonne Universite, CNRS, INSERM, 75006 Paris, France

## Abstract

Paul Green hypothesized that growth anisotropy of plant cylindrical organs could be controlled by cell-wall elastic strain. The present study aimed to challenge this hypothesis through a robust experimental and analytical framework. By combining live-cell imaging of *C. corallina* internodal cells with controlled turgor pressure manipulation, we simultaneously measured, for the first time, both the growth strain rate tensor and the elastic compliance tensor derived from multiaxial mechanical testing in the same cell. Under Green’s hypothesis, a significant correlation should be observed between the two tensors in all directions. Our results revealed a moderate yet significant correlation between multiaxial elastic compliances and growth strain rates most pronounced in the axial direction. The ratio of axial-to-radial elastic compliance was significantly correlated with the ratio of radial-to-axial growth strain rates. In contrast, other quantities, such as the radial compliance components or the orientations of the two tensors relative to the cell axis showed no significant correlation. Furthermore the growth strain rate tensor was strongly age-dependent in both magnitude and orientation, unlike the elastic compliance. Finally, analysis of intra-tensor variability revealed that axial and radial components were strongly correlated for both tensors, with a lowered correlation in the principal axis decomposition.

## 1 Introduction

Cylindrical organs play a central role in the plant world, encompassing stems, roots, root hairs, and trunks. The earliest terrestrial plants consisted of simple, ramified, photosynthetic stems, from which more complex organs—such as leaves and flowers—later evolved [1]. Growth of these cylindrical structures allows plants to adapt to their environment despite their immobility by efficiently coordinating two complementary processes: axial elongation, which enables exploration and optimizes the capture of resources (light, water, nutrients), and transverse thickening, which provides mechanical stability and prevents buckling. Understanding the regulatory mechanisms that control this anisotropic growth is therefore of broad relevance, as most plant organs originate from stem-derived structures and many of the underlying regulatory pathways have been conserved throughout evolution. The coordination of growth in different directions integrates a diversity of signals, from internal hormonal cues to environmental stimuli such as light, gravity, and mechanical forces [2].

The internal pressure supported by the plant cell wall is comparable to that borne by a car tire, and the resulting wall tension constitutes a major mechanical stimulus [3]. Building on three key observations made in the 1950s and 1960s on giant cylindrical *Characean* cells, Paul Green hypothesized that this stimulus might play a central role in regulating anisotropic cell wall expansion [4, 5].

Green first reported in 1955 that mature internodal cell growth follows an allometric, temperature-sensitive law: at 22 °C (resp. 36 °C), the longitudinal growth rate is 5 (resp. 2.5) times higher than the transverse growth rate [6]. He later observed in 1958 that the cell wall is reinforced by microfibrils oriented predominantly perpendicular to the cylinder axis [7]. These two findings were subsequently confirmed by Preston and colleagues [8], who added a third important observation: uniaxial mechanical tests on *N. axilaris* cell walls revealed an approximately 5:1 ratio between axial and radial elastic compliances in the fastest-growing cells (axial growth rate *>* 10% per day), and a reduced 2:1 ratio in the slowest-growing cells (axial growth rate < 5% per day) [9] . For the fastest-growing cells, the high compliance ratio, together with his earlier observations, led Green to suggest that anisotropic cell wall expansion is governed by a greater facility for separating microfibrils perpendicular to their orientation [3, 4]. A final line of evidence strengthened this view: randomization of microfibril deposition through herbicide treatment leads to isotropic growth [4, 10, 11].

Nonetheless, many questions remain regarding how, and to what extent, elastic compliances and growth rates are interconnected. Key information—such as temperature—is missing from the study by Probine and Preston [9], which makes direct comparison with the very limited measurements of axial–radial growth rates available in the literature difficult (only two measurements in [6] and one in [8]). It also remains unclear why Green’s hypothesis would apply only to the fastest-growing cells reported by Probine and Preston [9] but not to the slowest-growing ones. Measurements of the elastic properties of *Nitella* cell walls have also proven to be variable across subsequent studies (see tables in Supplementary Material §1; [9, 11, 12, 13]). Furthermore, in vivo plant cell walls experience multiaxial stresses, raising the question of how elastic constants derived from uniaxial tests [9] relate to the stresses actually sensed during growth. Multiaxial tests performed by Kamiya and collaborators [14] show that radial and axial strains behavior is non-linear as pressure goes from 0 to full turgor—whereas the uniaxial tests mentioned above were clearly conducted outside the linear regime [9]. Another open question concerns the thermosensitivity of growth anisotropy: If Green’s hypothesis is correct, how can the anisotropic growth ratio be strongly temperature-dependent [6] when elastic compliances appear to be much less affected by temperature (as suggested by Figure 8 of [15])? Finally, Green’s mechanistic explanation implicitly assumes an instantaneous regulation of growth by elastic strains, whereas the experiments supporting his allometric growth law span several days—far longer than the timescale of a transcriptional response (∼30 min). This discrepancy leaves room for alternative interpretations of the underlying regulatory processes.

Despite all these questions, Green’s hypothesis has experienced a recent revival in its multiaxial version thanks to an elegant experiment showing a nice collapse between multiaxial elastic strain anisotropy and growth strain anisotropy in the apical region of lily pollen tubes. Yet, this study makes the strong and unverified assumption that elastic moduli are isotropic and constant in this apical region [16]. This observation aroused some hope and prompted theoretical attempts to predict growth anisotropy from multiaxial elastic strain anisotropy [17, 18].

Understanding whether growth anisotropy is quantitatively controlled by cell-wall elastic strain is a key open question, and it is the central focus of this study. Addressing it requires the simultaneous measurement of stresses, multiaxial elastic strains, and growth strains with minimal noise, and in conditions that allow direct quantitative comparison. *Characean* internodal cells offer an ideal system for this purpose thanks to their simple cylindrical geometry, which inherently imposes a fixed 1:2 stress ratio between the longitudinal and circumferential directions. *Characean* internodal cell is also an interesting system as the biochemistry underlying cell wall growth is simpler than for higher plants whose growth mechanisms is mediated by enzymes: *C. corallina* relies on enzymeless chemistry to control both the axial rate of wall enlargement and the deposition of pectate into the wall [19]. However, measuring multiaxial elastic and growth strains on timescales shorter than the transcriptional response (∼30 min) is technically challenging, because the associated displacements are only a few times larger than the optical diffraction limit. Traditionally, growth has been tracked using latex aminated beads as fiducial markers adhered to the cell wall [16, 20]. Provided that the number of beads is sufficiently large and that they are well distributed, this technique can resolve displacements smaller than the optical resolution. The associated image-analysis task corresponds to a classical computer vision problem [21], closely related to “point-based registration methods for anisotropic scaling” [22].

This study aims to statistically challenge Green’s hypothesis through a robust experimental and analytical framework. We perform two successive experiments: a first experiment at 30 °C to measure the full growth strain rate tensor (Figure 1a_1_), immediately followed by a second, growth-inhibited experiment at 9 °C (Figure 1b_1_) that combines microscopy with the pressure-probe technique to determine the full tangent elastic-compliance tensor at full turgor. Two parametric models are used to estimate the tensors. The first one divides the angular span in four sectors and performs a simple directional average; it makes no assumption on the displacement fields and provides a baseline estimate of the tensors. The second parametric model is a first order Taylor expansion of the first fundamental form (a differential geometry tool for strain measurement [23]). This model assumes regular displacement fields and yields both the growth strain rate tensor and the elastic compliance tensor. This approach enables, for the first time, a direct quantitative comparison between the two tensors. According to Green’s hypothesis, a significant correlation between their components should be observed in all directions. Our approach provides a rigorous statistical framework to test this prediction.

**Figure 1:**
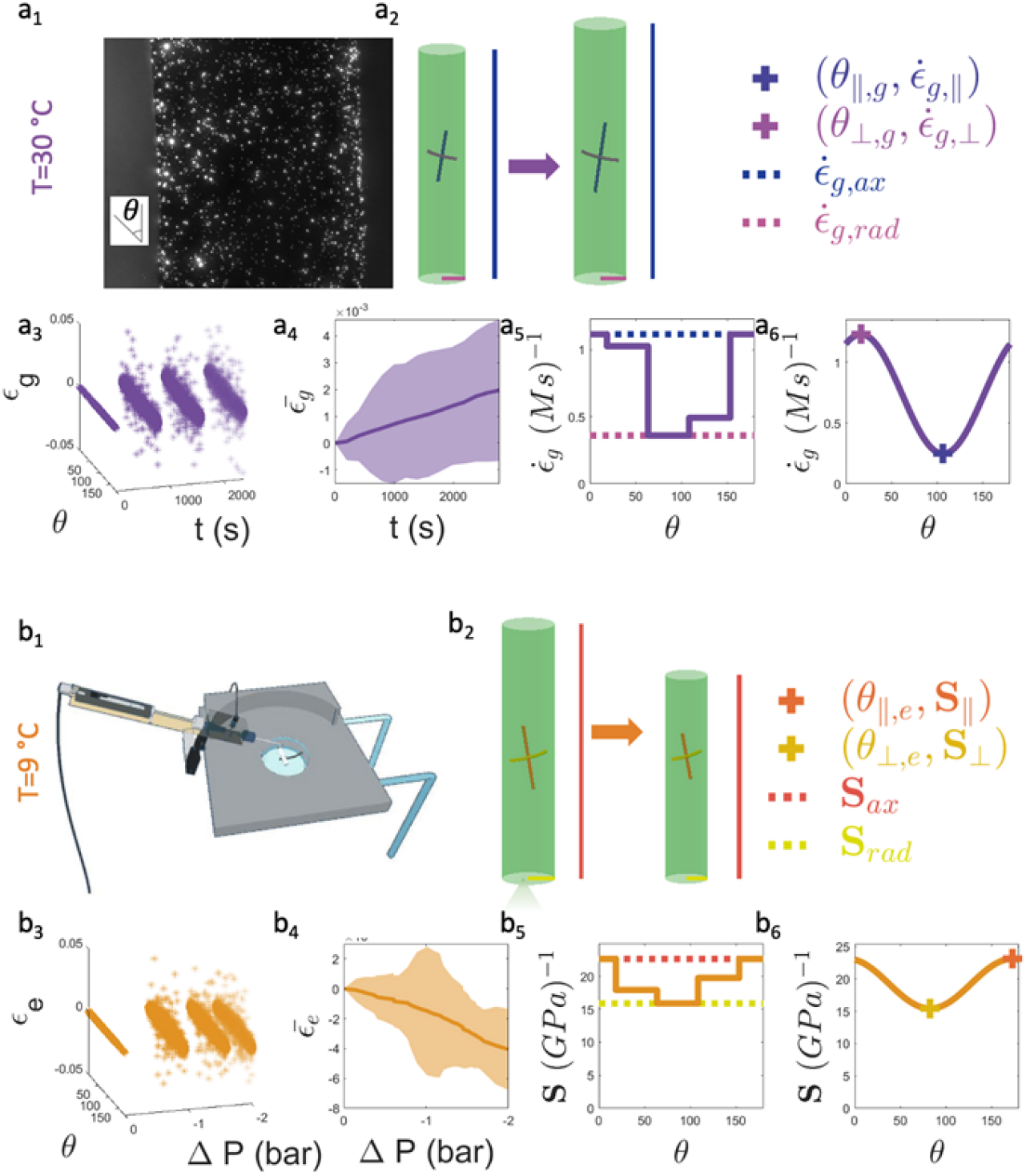
**a**_**1**_**–a**_**6**_. **Growth experiment at 30 °C. a**_**1**_. *C. corallina* internodal cell coated with latex a minated beads used as fiduciary markers (20x objective). **a**_**2**_. Left (resp. right): schematic of an internodal cell at the beginning (resp. end) of the experiment. The axial (resp. radial) direction is indicated by the vertical (resp. horizontal) light magenta (resp. light blue) line. The principal direction ∥ (resp. ⊥) is indicated by the tilted magenta (resp. blue) line in the middle of the cell. **a**_**3**_. Growth strain function of angle *θ* and time *t*. **a**_**4**_. Average growth strain 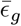 as a function of time *t*. The shaded area represents the standard deviation around the mean. **a**_**5**_. Estimated 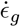 from the first parametric model. The magenta (resp. blue) dashed line corresponds to 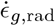 (resp. 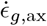). **a**_**6**_. Estimated 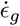 from the second parametric model. The dark magenta (resp. dark blue) cross marks 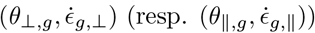. **b**_**1**_**–b**_**6**_. **Elastic experiment at 9 °C. b**_**1**_. Diagram of the pressure probe mounted on an epifluorescence microscope (20x objective). **b**_**2**_. Left (resp. right): schematic of an internodal cell before (resp. after) turgor manipulation. The axial (resp. radial) direction is indicated by the vertical (resp. horizontal) yellow (resp. red) line. The principal direction ∥ (resp. ⊥)is indicated by the tilted dark yellow (resp. orange) line in the middle of the cell. **b**_**3**_.. Elastic strain function of angle *θ* and time *t*. **b**_**4**_.. Average elastic strain 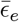 as a function of time *t*. The shaded area represents the standard deviation around the mean. **b**_**5**_. Estimated **S** from the first parametric model. The yellow (resp. red) dashed line corresponds to **S**_*rad*_ (resp. **S**_*ax*_). **b**_**6**_. Estimated **S** from the second parametric model. The dark yellow (resp. orange) cross marks (*θ*_⊥,*e*_, **S**_⊥_) (resp. (*θ*_∥,*e*_, **S**_∥_)).

## 2 Methods

### 2.1 Plant material and growth conditions

*Chara corallina* was cultivated in large, aerated tanks containing 100 L of nutrient solution following the protocol of Christopher Plieth (available on request). One day before the experiment, internodal cells were carefully selected and were left overnight in artificial pond water medium (APW; composition in mol/m3: NaCl 1.0, KCl 0.1, CaCl_2_ 0.1, and MgCl_2_ 0.1). The selected cells varied in size, exhibiting radii between 100 μm and 500 μm and lengths ranging from 150 μm to 1.5 cm. Prior to experimental manipulation, cells were gently cleaned off adherent leaves and debris under a binocular microscope and then thoroughly rinsed with fresh culture medium to remove any external contaminants.

### 2.2 Growth experiment

#### 2.2.1 Sample preparation and fluorescent labeling

Samples were prepared as either single internodal cells or pairs of naturally connected internodes to explore mechanical and growth anisotropy under conditions closely mimicking the natural physiological state. Paired internodes were carefully dissected while maintaining their natural connections intact to preserve intercellular mechanical continuity. 1 μm latex amine-modified fluorescent beads (L1030 Sigma-Aldrich) were used as fiduciary marker to track cell surface deformation. Beads were gently and uniformly attached to the cell wall surface by brief, gentle vortexing in an Eppendorf tube containing 1 μL of beads (Figure 1a_1_).

Samples (*N* = 36) were secured onto 20 mm Petri dishes using thin adhesive tape applied to the basal region of the larger cell, allowing both basal and apical cells to be studied simultaneously. Care was taken to avoid compressing the cell walls during mounting, ensuring that natural mechanical and growth behaviors were maintained. Additional adhesive applied to the tape edges prevented detachment during prolonged imaging sessions, facilitating stable measurements over extended experimental duration.

#### 2.2.2 Imaging system

The imaging system employed a Hamamatsu ORCA-Flash4.0 v3 camera mounted on an Olympus BX51 microscope. We use a CoolLED pE-4000 illumination source for fluorescence excitation. Precise spatial control during imaging was achieved through a *xyz* motorized stage with sub microns resolution (Märzhäuser mfd 5x + scan 75 × 50), facilitating accurate and reproducible multi-position imaging. All equipments were controlled and synchronised with a custom java software using micromanager drivers.

#### 2.2.4 Data acquisition

Time-lapse 3D fluorescence imaging was conducted at intervals of 1–2 min using 488 nm excitation. For cells larger than the microscope field of view (600 μm by 600 μm), imaging was limited to the central region to maintain consistent resolution and quality. The piezoelectric stage facilitated precise repositioning for capturing multiple regions along the cell. Imaging parameters were carefully optimized to balance minimizing photo-bleaching with sufficient temporal resolution for accurate tracking of cell deformation and growth. Additionally, a supplementary 0.1 W LED light source was positioned at the cell terminus to mimic natural lighting condition and providing lighting in-between the stack acquisition.

#### 2.2.4 Temperature control

The existing literature on axial growth is abundant with very good temporal resolution [15, 24] (minute time scale), whereas literature on radial growth is much scarcer with a low temporal resolution (a few days) [6, 8]. The difficulty lies in the much lower displacement associated to radial growth. To make the temporal resolution as fine as possible, we chose to conduct cell growth experiments at 30 °C to promote growth and maximize displacement.

Temperature control was critical to the experimental procedure and was achieved using dual interconnected LAUDA RE 415 cooling thermostats. This setup permitted precise and independent regulation of both the microscope stage and the culture medium temperatures.

#### 2.2.5 Growth duration

Growth experiments lasted between 20 min and 1½ h. The shortest acquisition duration corresponded to the largest cells: the field of view situated at the middle of the internodal cell tended to go out of focus quickly due to the growth of the cell part between the field of view and the anchorage point. The 30 °C temperature of the growth measurement was chosen to maximize the growth displacement relative to microscopic noise during this lag; at the end of each experiment growth strains were of the order of a few tenths of percent.

#### 2.2.6 Tracking of fiduciary marker trajectory

The field of view observed with a 20x (Olympus UPlanFL N 20x / 0.50 ∞ / 0.17 / FN 26.5 UIS 2) objective was a part of the cell situated at the center of the internode. The first step of the analysis was a rigid registration (Registration ImageJ plugin) to remove the bulk movement due to the growth or elastic deflation of the cell situated between the anchorage and the field of view. The second step was to retrieve the beads trajectories from 2D projections analyzed using the ImageJ plugin Trackmate [25]. Outlier trajectories due to beads floating in the medium were detected by Trackmate and removed from further analysis. The third step was to get accurate *z*-coordinate determination of bead positions through Gaussian fitting of intensity profiles in the full 3D image stacks.

The localization precision [26, 27] has been estimated observing the standard deviation in 85 repeated acquisition stack of latex aminated beads position adhering on an inox thread (diameter of 150 μm, similar to the smaller internodal cells) maintained at constant 30 °C temperature to avoid thermal dilatation or retraction during a 43 min acquisition time (typical duration of the experiment). The average (on all the beads) standard deviations were 83 nm in *x*, 93 nm in *y*, and 379 nm in *z* (see Figure 2 in Supplementary Material §2).

**Figure 2:**
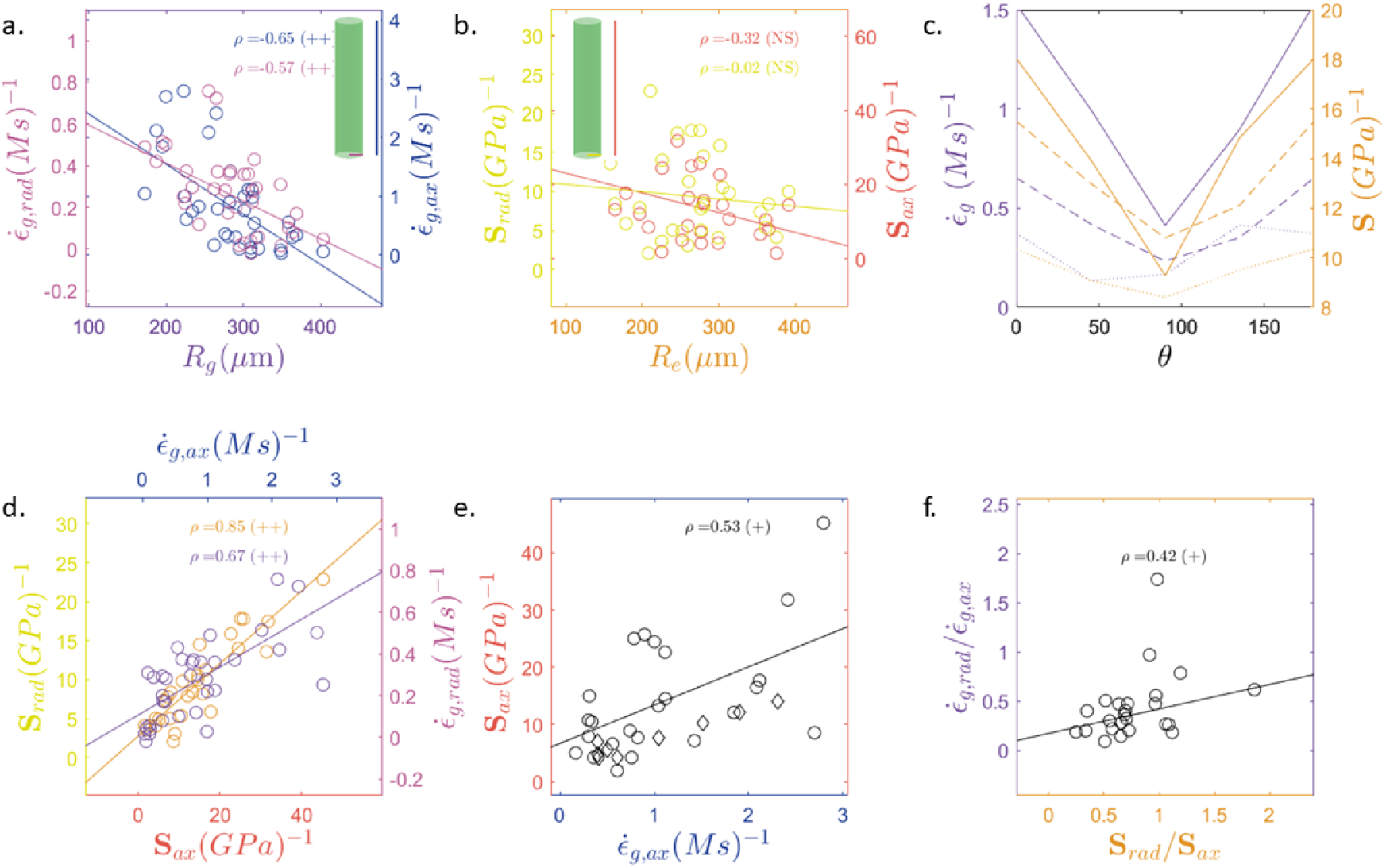
**a**. Radial growth strain rate (magenta) and axial growth strain rate (blue) vs. cell radius *R*_*g*_ (*N* = 35). **b**. Radial elastic compliance (yellow) and axial elastic compliance (red) vs. cell radius *R*_*e*_ (*N* = 28). **c**. Data pooled into three quantiles based on cell radius (*N* = 24). The first quantile is shown as a solid line, the second as a dashed line, and the third as a dotted line (*N* = 28 for elastic compliance, *N* = 35 for growth strain rate). Right axis (orange): average angular distribution of elastic compliance for each quantile. Left axis (violet): average angular distribution of growth strain rate for each quantile. **d**. Left axis (orange): radial elastic compliance versus axial elastic compliance (*N* = 28). Right axis (violet): radial growth strain rate versus axial growth strain rate (*N* = 35). **e**. Axial elastic compliance versus axial growth strain rate (*N* = 24). (o) Our Data. (⋄) Data extracted from [15]: The growth velocity reported in this reference was converted into a growth strain rate by assuming an internode length of 1 cm, as stated elsewhere in the same reference. *ρ*: Spearman correlation coefficient. **f**. Ratio of growth strains versus ratio of elastic compliances (*N* = 24). In all panels, regression lines were computed using MATLAB’s `fitlm` function with the robust `‘bisquare’` option. NS: *p*-value ≥0.05; +: *p*-value < 0.05; ++: *p*-value < 0.001. The growth strains correspond to the (CD) condition. The corresponding figure for the (SD) condition is Figure 3 in the Supplementary Material.

From the *N* = 36 samples, one of the cells was excluded from the growth analysis because the number of beads at the end of the acquisition was too low (*N*_beads_ < 50), as the cell went out of focus.

#### 2.2.7 Growth strain rate estimation

##### Alignment procedure

The shape of *Chara corallina* internodal cell is approximately cylindrical, but the principal strain axes are not necessarily aligned with this cylinder, as suggested by the spiral behavior of growing cells [28] (Figure 1a_2_). To test this hypothesis, the first step of the analysis consisted of determining the principal axis and radius of the cell by observing the *xz* and *yz* projections of the point cloud. As internodal cells are cylindrical, the cylinder boundaries could be well approximated by straight lines, which were manually identified on the two projections. The point cloud was then aligned so that these boundaries were parallel to the *x* axis in both projections. A second correction step consisted of dividing the cylinder into five segments along the *z* axis. For each segment, a circle was fitted to the corresponding *xy* coordinates. The principal direction of the five circle centers was then determined, and the point cloud was rotated so that this principal axis aligned with the *z* direction. At the end of the procedure, the cell radius *R*_*g*_ was estimated by fitting a circle to the *xy* coordinates of the rotated point cloud.

##### Isometric mapping of the cylindrical cell on a plane

The second step simplified strain estimation by mapping the point cloud located on the cylinder surface onto a plane, while preserving both the angular coordinate around the cell axis, *θ*, and the bead–bead geodesic distances *L*. This was done using the following isometric transformation:

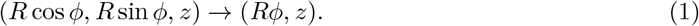

(*R, ϕ, z*) being the coordinate of the cylinder surface in a cylindrical coordinate system with the same axis as the cylinder. *ϕ* belongs to [−*π, π*]. For the *i*^th^ beads pair constituted by the beads *i*_1_ and *i*_2_, the distance between the two beads at time *t* was given by:

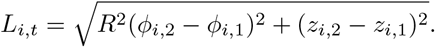

The inclination respective to the cell axis of the segment joining one bead from the other *θ*_*i,t*_ was provided by the four-quadrant inverse tangent (in degrees) of (*z*_*i*,2_− *z*_*i*,1_) and (*Rϕ*_*i*,2_ − *Rϕ*_*i*,1_) modulo 180°.

##### Growth strain determination

For each experiment, the number of beads adhering to the cell ranged between 55 and 1500, which corresponded to a number of pair of beads, *N*_pairs_ ranging between 1500 and 10^6^. The growth strain *ϵ*_*g,i*_(*t*) associated to the *i*^th^ pair of beads over the time window [0, *t*] is provided by:

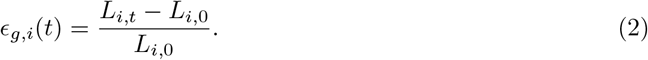

Because growth-induced displacements were of the same order of magnitude as the localization error directly applying this formula leaded to a highly noisy output (Figure 1a_3_). As a consequence, the growth tendency could not be reliably detected at the level of individual bead pairs. However, thanks to the large number of bead pairs, the averaged growth strain was well above the noise level and could be robustly measured (Figure 1a_4_; Figure 2 of Supplementary Material §3.2).

The average growth strain over the pairs of beads was estimated at each growth step *t* by estimatingthe parameters 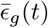 of the following linear model with `fitlm` of MATLAB:

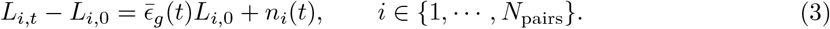

where *n*_*i*_(*t*) is noise. This linear model reduced the impact of localization error compared to directly averaging growth strains. Indeed, because the growth-strain formula contains *L*_*i*,0_ in the denominator, bead pairs with very small initial spacing would disproportionately amplify noise if growth strain were directly averaged (see Figure 2 of Supplementary Material §3.2). This step was used later to determine the 20 min of maximal average growth strain (see below).

##### Direction-specific growth strain rates

We aimed to determine the direction-specific growth strain rates. Therefore, the angular domain [0, 180] of the variable *θ*_*i*,0_ was divided into four angular sectors:

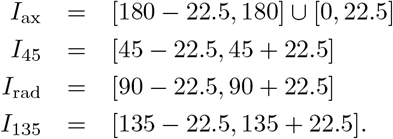

The growth strain rate is defined by:

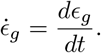

To quantify the influence of the growth direction on this quantity, we considered the temporally and spatially averaged direction-specific growth strain rates 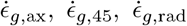 and 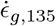. These parameters were estimated using MATLAB’s `fitlm` with a bisquare robust option applied to the linear regression model (see Figure 1a_5_):

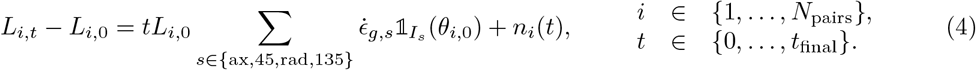

##### Principal growth strains rate

The growth strain rate tensor coefficient *δE*_*g*_, *δF*_*g*_, *δG*_*g*_ were obtained by fitting the parameters of the following linear model using the same bisquare-robust procedure:

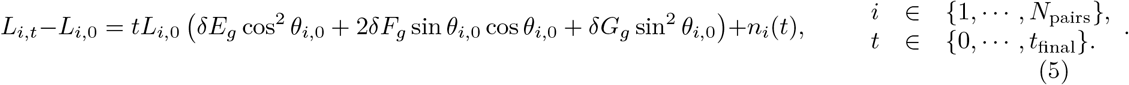

The principal axes were obtained by diagonalizing the tensor:

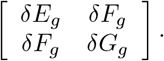

We denote by *θ*_∥,*g*_ (resp. *θ*_⊥,*g*_) the eigen-direction closest to the cell axis (resp. radial axis), and 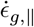 (resp. 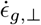) the associated eigenvalue (Figure 1a_6_ and Annex 3.2). According to the *p*-values returned by fitlm, the coefficients *δE*_*g*_, and *δG*_*g*_ all differed significantly from zero (*p* < 10^−7^) and the coefficients *δF*_*g*_ all differed significantly from zero (*p* < 10^−7^) except for one cell (N=35).

##### Analysis on shorter 20 min duration windows

Experimental constrains made the duration of the different acquisitions ranging from 20 min to 90 min which would not have influenced the results if all the cell were in stationary growth regime. Though the stationary assumption holds for the majority of the 35 cells; some of them did not grow throughout the entire experiment and even exhibited transient retraction phases. As our study aims to understand how grow strains and elastic strain are related, the rate coefficients were computed twice with two choice of temporal windows:

- over the complete duration (CD)
- over a shorter interval (SD) corresponding to the maximal average growth strain increment over a 20 min duration (see red boxes in Figure 2, Supplementary Material §3.2).

### 2.3 Multi-axial mechanical test

#### 2.3.1 Pressure probe implementation

The pressure measurement system was based on the design previously described by [29], consisting of a custom-designed plexiglass chamber with a T-shaped 1 mm channel (Figure 1b1) (manufactured by Der Werkzeugmacher, Eckersdorf Germany). One end of the channel was connected to a glass capillary via a plexiglass nut, while the opposite end housed a metal piston. Piston control was automated by motorizing the micromanipulator using a stepper motor driven by an Arduino controller, allowing precise pressure adjustments without manual intervention. This modification provided feedback control of the piston position and, consequently, more stable pressure regulation throughout experiments. A Honeywell 26PCGFA6D pressure sensor connected to a Burster 9236 amplifier provided continuous pressure measurements. Overall, this automated system significantly improved the stability and reproducibility of pressure measurements compared to manual operation, while maintaining the proven fundamental design principles of the original pressure probe system.

The pressure probe was mounted directly onto the microscope *x*–*y*–*z* translation stage using a Narishige MN-153 micromanipulator fixed to the cylindrical axis of an angle-transmission reduction gearing system (Prudhomme Transmission, Model RVP type 2, combination E, ratio 60:1), allowing fine control of the probe inclination relative to the sample plane (Figure 1b_1_). The capillary tip was beveled at a 30° angle, yielding an aperture between 20 μm and 50 μm depending on the radius of the cell to be attached. Sticking operations were performed under a 2.5x objective. The internodal cell was attached by passing the probe through the nodal cell. Pressurized cytoplasm first entered the capillary, which was pre-filled with low viscosity silicone oil (Wacker AK 0.65); the meniscus was then brought back to the capillary tip to obtain an accurate pressure measurement.

#### 2.3.2 Pressure perturbation protocol

Multi-axial mechanical tests were performed at a temperature of 9 °C using a LAUDA RE 415 temperature-control system. This temperature reduction was essential for minimizing growth during mechanical measurements and for obtaining more reproducible elastic strain estimates [15]. Nevertheless, a low temperature also enhances cytoplasm viscosity which could to non trivial behavior. After temperature stabilization, cells were punctured with the pressure-probe micro-capillary. Two types of pressure perturbations were analyzed depending on the cell’s response to puncture:

- **Natural pressure decreases**: In some cells, slow leaks developed after probe insertion. In these cases, we recorded the gradual pressure decrease while simultaneously imaging cell deformation. These natural pressure drops provided controlled deformation data without additional manipulation. The fluid velocity associated with the leak was very low (< 2 mm/h). Using Poiseuille’s law for a cylindrical channel connecting the sensor to the meniscus (diameter 1 mm, length 10 cm) filled with the silicon oil (viscosity 5 × 10^−4^ Pa.s at a 25 °C temperature), this flow rate corresponded to a relative error of less than 10^−5^ in the pressure measurements.
- **Imposed pressure steps**: For cells that maintained a stable turgor after puncture, we applied controlled pressure steps using the motorized piston. The magnitude of these steps was chosen to match the typical pressure variations observed in leaking cells.

For both types of perturbations, cell deformation was monitored using 3D bead tracking. The resulting deformation fields were subsequently analyzed using the angular strain analysis described in the following section.

Reference [14] clearly shows that the cell wall operates in a non-linear elastic regime when pressure varies from 0 to full turgor. For this reason, pressure manipulations in our experiments were limited to a maximum of 1 bar around full turgor, in order to measure the tangent elastic compliances within the local linear regime. The complete sequence of cooling the bath and puncturing the cells required between 30 min and 1 h, depending on the time needed to position the pressure probe, attach the cell, and perform the meniscus adjustments. The chosen field of view was 600 μm. At the end of each acquisition, the metal piston was moved and the connection with the cell was checked by observing the corresponding movement of the meniscus under the 2.5x objective. Out of the *N* = 36 samples, 8 cells were excluded from the elastic analysis because the capillary was plugged for a total of *N* = 28 valid samples.

#### 2.3.3 Elastic strain and elastic compliance estimation

##### Alignment procedure

The alignment procedure was repeated. Because the portions of the cell observed under the reduced field of view provided by the 20x objective did not necessarily correspond from the growth experiment to the pressure manipulation experiment. The relative difference 2 |*R*_*e*_−*R*_*g*_| */*(*R*_*e*_ +*R*_*g*_) between the radii *R*_*e*_ obtained from the cylindrical fit of the pressure manipulation data and the radius *R*_*g*_, corresponding to the growth data could be as high as 30%. Cells whose estimated radii differed by more than 11% between the two successive experiments were excluded from the comparison analysis with growth strain rates (*N* = 24) but maintained for the purely elastic analysis (*N* = 28).

##### Elastic strain determination

The elastic strain *ϵ*_*e,i*_(*t*) associated to the *i*^th^ pair of beads over the time window [0, *t*] was defined by:

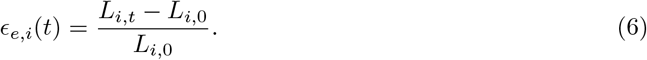

At each time *t*, elastic strains were determined by estimating the parameters of the same linear model as for growth strain but for the data obtain during the multi-axial mechanical tests (see Supplementary Material §3.2). The elastic strain were of the order of a few tenths of percent for a pressure manipulation of 1 bar.

##### Angular dependence of the elastic compliances

The elastic compliance was defined:

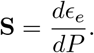

(**S**_ax_, **S**_45_, **S**_rad_, **S**_135_), the elastic compliances in the four directions *θ* = 0°, 45°, 90°, 135° were estimated from the pressure-perturbation data using the same procedure as for the growth strain rate, except that the variable *t* was replaced by the pressure increment *P* (*t*) − *P* (0) in Equation (5) (Figure 1b_5_).

##### Principal elastic compliances

The coefficients *δE*_*e*_, *δF*_*e*_, *δG*_*e*_, and the principal directions *θ*∥_,*e*_ (resp. *θ*⊥_,*e*_), the closest to the cell axis and (resp. closest to the radial direction), together with their associated compliances **S**_∥_ and **S**_⊥_ were retrieved using the same model as for principal growth-strain rates, again replacing *t* by *P* (*t*) − *P* (0) in Equation (4) (Figure 1b_6_ and Supplementary Material §3.2).

According to the *p*-values returned by `fitlm`, the coefficients *δE*_*e*_, and *δG*_*e*_ all differed significantly from zero (*p* < 10^−4^) and the coefficients *δF*_*e*_ all differed significantly from zero (*p* < 10^−4^) except for one cell (N=28). Some of the cells showed negative compliance in some directions (radial or axial) which could be explained by the high viscosity of the cytoplasm at low temperature.

### 2.4 Correlation analysis

In all the figures presented in this article, regression lines were computed using MATLAB’s `fitlm` function with the option `‘robust’` and the `‘bisquare’` algorithm. Each correlation analysis was first performed using the complete-duration (CD) growth strain rate coefficients and then repeated using the reduced-duration (SD) coefficients. Because the (CD) and (SD) correlation coefficients differed only marginally, the discussion in the main text focuses primarily on the (CD) coefficients. Both (CD) and (SD) correlation coefficients and their associated *p*-values are provided in the Supplementary Material (§4.1 and §4.2; Tables 1–8).

## 3 Results and discussion

### 3.1 Association of growth and elastic strain in cylindrical principal coordinate

#### Correlation of cellular properties with age

In this study, the cell radius was used as a proxy for the cell age, as the growth protocol in a 100 L water tank made it difficult to evaluate the exact age of *C. corallina* internodal cells. The length of the 35 cells examined were all below 15 mm due to the spatial limitation imposed by the LAUDA thermalization system. Axial and radial growth strain rates were significantly and negatively correlated with radius (Spearman *ρ* = −0.65, *p* = 3×10^−5^ and *ρ* = −0.57, *p* = 5×10^−4^, respectively) (Figure 2a), as were the coefficients 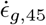 and 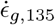 (*ρ* = −0.71, *p* = 5×10^−6^ and *ρ* = −0.45, *p* = 0.007 respectively) (see Supplementary Material §4.1). The tendency of axial and radial growth strain rates to decrease with age had already been reported in the limited literature data (*N* = 2, Figure 5 in [6]; *N* = 1, Figure 3 in [8]) but it is the first evidence for the shear components.

**Figure 3:**
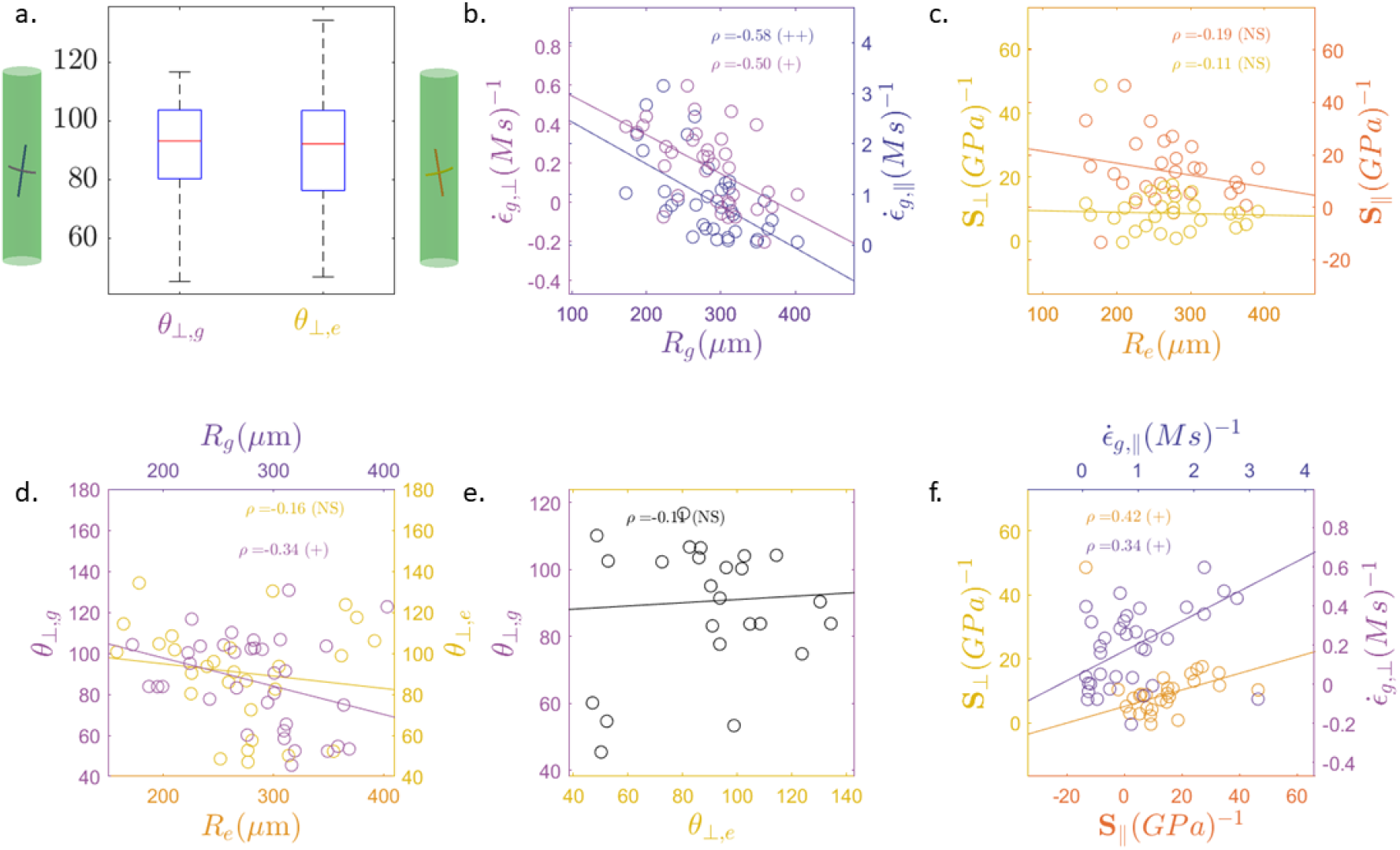
**a**. Boxplots of the principal axis direction of the elastic compliance tensor ⊥ component (light orange) and the growth strain rate tensor ⊥ component (light violet) (*N* = 24). **b**. The principal growth strain rate in the direction (light violet) and in the ∥ direction (dark violet) vs. cell radius *R*_*g*_ (*N* = 35). **c**. The principal elastic compliance (yellow) in the ⊥ direction (light orange) and in the ∥ direction (dark orange) vs. cell radius *R*_*e*_ (*N* = 28). **d**. The principal growth strain rate direction (respectively, elastic compliance) in the ⊥ direction (light violet, respectively light orange) and in the ∥ direction direction (dark violet, respectively dark orange) vs. cell radius *R*_*g*_ (respectively *R*_*e*_) (*N* = 35, respectively *N* = 28). **e**. Principal growth strain rate in ⊥ the direction (light violet) vs. principal elastic compliance in the ⊥ direction (light orange) (*N* = 24). **f**. (Left axis, orange) Elastic compliance in the ∥ direction vs. elastic compliance in the ⊥ direction (*N* = 28). (Right axis, violet) Growth strain rate in the ∥ direction vs. growth strain rate in the ⊥ direction (*N* = 35). In all panels, regression lines were calculated using MATLAB’s `fitlm` function with the robust `‘bisquare’` option. *ρ* is the Spearman correlation coefficient. NS: *p*-value ≥0.05; +: *p*-value < 0.05; ++: *p*-value < 0.001. The growth strains correspond to the (CD) condition.The corresponding figure for the (SD) condition is Figure 4 in the Supplementary Material.

To characterize how the angular distribution of the growth strain rate (relative to the cylinder axis) evolved with age, we divided the sample into three quantiles based on cell radius (*R* < 265 μm, 265 μm ≤*R* < 309 μm, 309 μm ≤ *R*). For each quantile, the growth strain rate coefficients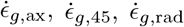 and 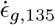 were averaged (Figure 2c). For the first quantile (small radii), the growth strain rate distribution showed a clear minimum in the radial direction, whereas for larger cells it became more uniform. The tendency of growth strain rate to decrease with radius could also be observed.

The four components of the elastic compliance (axial **S**_ax_, **S**_rad_ and the two shear terms **S**_45_, **S**_135_) were negatively correlated with the radius but none of the correlations was significant (*ρ* = −0.32, *p* = 0.10; *ρ* = −0.30, *p* = 0.12; *ρ* = −0.02, *p* = 0.93; *ρ* = −0.23, *p* = 0.23) (Figure 2a). To explore this pattern further, we divided the cell sample into three quantiles based on radius (*R* < 245 μm, 245 μm ≤*R* < 283 μm, 283 μm < *R*). For each quantile, the elastic compliance coefficients (**S**_ax_, **S**_45_, **S**_rad_, **S**_135_) were averaged to obtain a representation of their angular distribution relative to the cylinder axis (Figure 2c). For the different quantiles, the elastic compliance distributions were similar, both showing a clear minimum near the radial direction; the tendency of the compliances to decrease with age could also be observed.

#### Correlation of cellular properties along the different cylindrical principal directions

A strong and significant correlation was observed for all of the 6 pairs of growth strain rate components in *C. corallina* (see Table 2 in Supplementary Material §4.1); indicating that variability in growth rates at constant temperature was not independent (Figure 2d). Similarly, correlations between pairs of elastic compliance components were also consistently significant (Figure 2d) (see Table 2 Supplementary Material §4.1).

The question of whether axial and radial growth in cylindrical organs were independently regulated has been examined in another experimental system, the *A. thaliana* root: differences in the temperature dependence of axial and radial growth led some authors to propose that these two directions may be regulated differently [30]. This observation, however, does not seem to apply to our system under constant temperature conditions.

#### Correlation of the mechanical and growth cellular properties

For the three radius quantiles, both the elastic compliance distribution and the growth strain rate distribution displayed a clear minimum along the radial direction (Figure 2c); the distribution tended to flatten with the radius. This qualitative similarity between the elastic compliance and growth strain rate distributions (Figure 2c) suggests that the two quantities may be related, in line with the hypothesis formulated by Paul Green [4, 12]: “The direction of yielding of a small area of cell surface must be a function of (1) the pattern of stresses within it and (2) its structure, particularly the alignment of reinforcing microfibrils.”

For each direction *θ* = 0°, *θ* = 45°, *θ* = 90° and *θ* = 135°, the correlation between the growth strain rate component and the corresponding elastic compliance component was positive (*ρ* = 0.53, 0.51, 0.03, 0.35) and significant for two components (*p* = 0.009, *p* = 0.01, *p* = 0.87, *p* = 0.1) (see Table 2) for the (CD) analysis. A Fisher combined probability test performed on the four components yielded a significant result (*p* = 3 × 10^−3^), indicating that growth strain rate component and elastic compliance were correlated overall. The significantly positive correlation for the axial component had already been reported by [15] for *C. corallina* cells growing at 23 °C corresponding to the oldest cells of our sample (∼1 cm long). The data points from [15] aligned well with our measurements (Figure 2e). This positive correlation between growth strain rate and elastic compliance was also in line with recent observation of the negative correlation between bulk modulus and growth rate for another experimental system *Marchantia polymorpha* [32].

The simplest function combining point (1) and point (2) of Green’s hypothesis (see above) is the ratio between axial and radial elastic compliances. Observations on *Lilium* pollen tube apical growth suggest a quantitative relationship between this ratio and the anisotropy of the growth strain rate (Figure 4 in [16]). In our analysis (Figure 2f), the correlation between elastic compliance anisotropy and growth strain rate anisotropy was at the threshold to be significant for both the (CD) analysis and the (SD) analysis (*ρ* = 0.42, *p* = 0.04 (CD); *ρ* = 0.41, *p* = 0.05 (SD)). The regression line slope was 0.25 for the (CD) analysis. Overall, our dataset supports extending the observation made in lily pollen tubes to *C. corallina* growing at 30 °C provided growth strain rate are estimated on long enough time scales.

### 3.2 Association between growth strain rate and elastic compliance in principal directions

#### Orientation of growth strain rates

For all the 35 cells in the study except one, *δF*_*g*_ was significantly different from 0 (*p* < 10^−7^), indicating that the principal direction of the growth strain tensor differed from the principal direction of the cylindrical geometry; it indicates the cell twists as it grows (Figure 3a). The distribution of principal directions was nearly symmetric and centered around a −3.1° inclination relative to the cylinder axis, with a standard deviation of 22.3° (Figure 3a).

The principal value of the growth strain rate tensor closest to the axial direction, 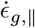 and 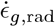 were strongly and negatively correlated with the radius (*ρ* = −0.50, *p* = 0.002; *ρ* = −0.58, *p* = 0.0004) (Figure 3b). The correlation between the principal direction of the growth strain tensor and the cell radius was negative and significantly different from zero (*ρ* = −0.34, *p* = 0.04)(Figure 3d). According to the regression line, the principal direction switches from dextral to sinistral around a radius of 270 μm. This tendency was already reported in the 1950s–1960s for a different but related system: the internodal cells of *N. axilaris* and other *Nitella sp*. [31, 28]. In the single experiment presented in Figure 4 of [31], Paul Green observed that the spiral orientation switched from dextral to sinistral approximately when the cell diameter reached a plateau—an observation also described by [28] when the cell reached a length of 12 mm.

#### Orientation of elastic compliances

For all the 28 cells in the study except one, *δF*_*e*_ was significantly different from zero (*p* < 10^−4^), indicating that the principal direction of the *C. corallina* elastic compliance tensor differed from the principal direction of the cylindrical geometry. In other words, the cell wall elastic law is non-orthotropic: the principal directions of the elastic tensor do not coincide with the cylinder’s principal axes. The distribution of these principal directions was symmetric. The average inclination was 1° relative to the cylinder axis, with a standard deviation of 25° (Figure 3a).

The correlation between the principal direction of the elastic compliance tensor and the radius was negative though not significantly different from 0 (*ρ* = −0.16, *p* = 0.43)(Figure 3d). The principal values of the elastic compliance tensor component were both negatively correlated with the radius but none of the correlations was significant (*ρ* = −0.11; *p* = 0.59; *ρ* = −0.19; *p* = 0.32; see Figure 3c).

The non-orthotropic nature of the cell wall was previously reported in *Nitella* species by Probine [28], who glued a small stick perpendicular to the axis of a *N. axillaris* internodal cell and subjected the cell to hyperosmotic shocks, observing that the stick rotated. Probine also described an age-dependent change in orientation, with the direction of stick rotation reversing when the cell length reached approximately 1.2 cm (Figure 2 in [28]). The trend is not significant in our experimental system involving *Chara corallina* internodal cells.

#### Correlation between elastic compliances principal direction and growth strain principal direction

According to [28], a cell length of 1.2 cm marks the point at which the spiral growth of *N. axillaris* shifts its orientation from dextral to sinistral. This suggests a strong correlation between the principal directions of the elastic compliance tensor and the growth strain rate tensor but no significant correlation was observed between the two principal directions (*ρ* = −0.11, *p* = 0.62 (CD), *ρ* = −0.23, *p* = −0.27 (SD); see Figure 3e). We did not test the correlation between the ratios of the principal values of 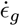 and **S** as the principal direction did not match.

#### Correlation of cellular properties along the different principal directions

The correlation between the two principal values of the growth strain rate tensor was lower than in cylindrical coordinates but was still significant (*ρ* = 0.34, *p* = 0.04 (CD); *ρ* = 0.44, *p* = 0.009 (SD); Figure 3f) as well as the correlation between axial and radial elastic compliances (*ρ* = 0.42, *p* = 0.03 (CD); *ρ* = 0.42, *p* = 0.03 (SD); Figure 3f).

## 4 Conclusion

The aim of this study is to challenge the sixty-year-old Green hypothesis for cylindrical *Characean* internodal cells and, more broadly, to clarify the relationship between growth strain and elastic compliance. To achieve this, we designed an experiment that provides, for the first time, the distribution along the cell axis of both multiaxial elastic compliances and growth strain rates, enabling statistical comparison between them.

Our results confirm earlier macroscopic observations in *Characean* internodes [31, 28]: both the elastic compliance tensor and the growth strain rate tensor exhibit significant shear components. We further refine the relationship between multiaxial elastic compliance and growth strain rates, showing evidence for a moderate yet significant correlation that is most pronounced in the axial direction. In contrast, the orientations of the two tensors relative to the cell axis showed no significant correlation. The ratio of axial to radial elastic compliance correlated significantly with the ratio of radial to axial growth strain rates, which is consistent with previous observations in lily pollen tubes [16]. However, in some cells, we observed that the ratio between growth strain rates varied substantially over short timescales (< 30 min), whereas the ratio between elastic compliances is expected to vary much more slowly. Therefore, we do not believe that the ratio between radial and axial elastic strains is a quantitative proxy for the relationship between radial and axial growth strain rates as hypothesized by some authors (including ourselves) [16, 17, 18].

The study also assesses how these distributions vary with cell radius, used here as a proxy for cell age. Our findings are consistent with earlier macroscopic observations showing that growth strain rates decline with age, accompanied by a shift in orientation from dextral to sinistral. By contrast, multiaxial elastic compliance appears far less age-dependent.

Finally, this work provides insight into how the different components of each tensor vary in relation to one another. For both elastic compliance and growth strain rate tensors, axial and radial components are significantly correlated. As a perspective, the robustness of this observation could be tested by introducing additional sources of variability, such as controlled changes in light or temperature.

## Acknowledgement

We thank Jacques Dumais, Ben Jordan, Christopher Plieth for sharing protocoles. Marie-Beatrice Bogeat Triboulot, Yoel Forterre and Geoffroy Guena for training EC to the pressure probe and François Heslot and Atef Asnacios for advices. We thank also Nadine Peyrieras and the BioEmergences platform at MSC, France BioImaging infrastructure ANR-10-INBS-04, ANR-11-EQPX-029.

## Funding

ANR JCJC ANR-20-CE30-0005.

## Notes

### Competing Interest Statement

The authors have declared no competing interest.

