## Supplementary Material for "Mechanical and Growth Anisotropy in *Chara corallina*: Challenging Green’s Hypothesis"

February 6, 2026

#### 1 Growth strain and elastic strain anisotropy measurements in the literature

The data in the three tables have been gathered and converted in ratio of strains, though some authors provide it as ratio of elastic moduli [8].

| Growth anisotropy | ($N = 2, T = 22$ °C) Fig. 5 [5] (*N. axilaris*) | ($N = 1, T = 36$ °C) Fig. 5 [5] (*N. axilaris*) | ($N = 1$) (IPC) Fig. 5[9] (*N. axilaris*) | ($N = 1$) Fig. 3 [7] (*Nitella sp.*) |
|---|---|---|---|---|
| $\dot{\epsilon}_{g,\text{ax}}/\dot{\epsilon}_{g,\text{rad}}$ | 5 | 2.25 | 0.2 | 5 |

| Elastic anisotropy (uniaxial test) | ($N = 12$) Fig. 3 [8] (*N. species*) | ($N > 5$) Fig. 2 [10] (*N. species*) | ($N > 5$) (IPC) Fig. 2 [10] (*N. species*) | ($N = 6$) Fig. 2 [12] (*C. corallina*) ($R =$ 0.5 mm) |
|---|---|---|---|---|
| $\mathbf{S}_{\text{ax}}/\mathbf{S}_{\text{rad}}$ | $[1.8 - 5.3]$ | $1.5 \pm 0.2$ | $0.8 \pm 0.1$ | $[2.2 - 2.9]$ |

| Elastic anisotropy (multiaxial test) | ($N > 5$) Fig. 6 [10] (*N. species*) | ($N > 5$) (IPC) Fig. 6 [10] (*N. species*) | ($N = 1$) Fig. 7 [13] (*Nitella*) |
|---|---|---|---|
| $\mathbf{S}_{\text{ax}}/\mathbf{S}_{\text{rad}}$ | $2.2 \pm 0.2$ | $0.8 \pm 0.1$ | 1.3 |

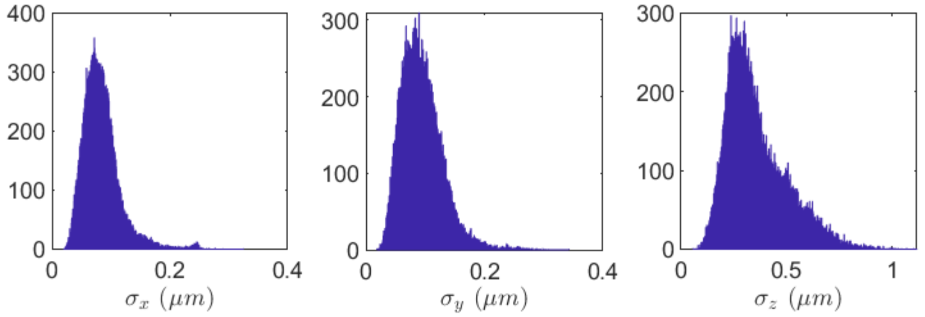


Figure 1: Histogram of the localization precision (standard deviation on 85 repeated acquisition stack) $\sigma_x$, $\sigma_y$ $\sigma_z$ for the $x$, $y$, $z$ the position of 65193 beads adhering on an inox wire thread maintained at constant temperature for 43 min.

### 2 Microscopic error

### 3 Cells

#### 3.1 Average growth strain

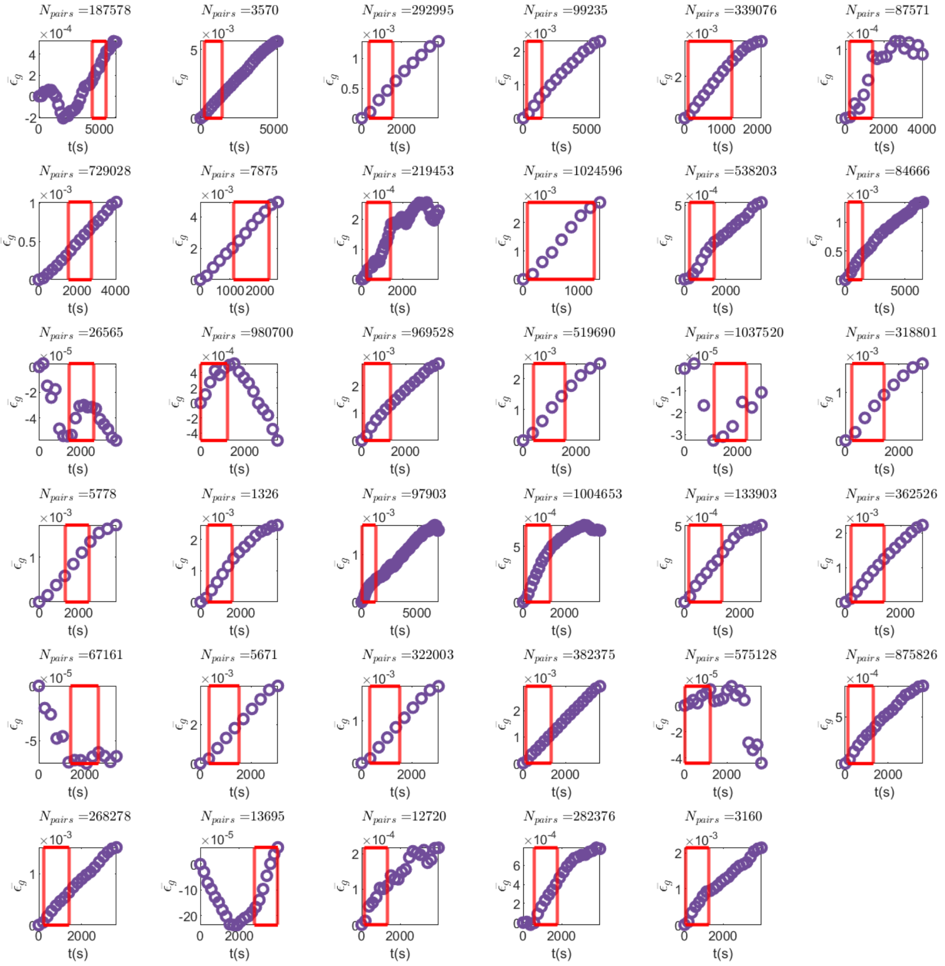

Figure 2: Average growth strain $\bar{\epsilon}_g$ vs. time $t$. The red box corresponds to the 20 min of maximal average growth strain increment.

#### 3.2 Growth strain and elastic strain

**Principal growth strains.** At each time point $t$ the principal growth strains were obtained by fitting the parameters $\delta E_{g,t}$, $\delta F_{g,t}$ and $\delta G_{g,t}$ of the model:

$$L_{i,t} - L_{i,0} = L_{i,0}\left(\delta E_{g,t}\cos^2\theta_{i,0} + 2\delta F_{g,t}\sin\theta_{i,0}\cos\theta_{i,0} + \delta G_{g,t}\sin^2\theta_{i,0}\right) + n_i(t), \quad i \in \{1, \cdots, N_{\text{pairs}}\}. \tag{1}$$

This formula is a first-order Taylor expansion of the first fundamental form (a differential geometry tool for strain measurement), valid for small length increments. This procedure yielded the components of the growth-strain tensor

$$\begin{bmatrix} \delta E_{g,t} & \delta F_{g,t} \\ \delta F_{g,t} & \delta G_{g,t} \end{bmatrix}.$$

We similarly estimated the components of the elastic-strain tensor: $\delta E_{e,t}, \delta F_{e,t}, \delta G_{e,t}$ by repeating the procedure on the elasticity data.

In the following panels, each row corresponds to one cell. For each row, the first panel (from left to right) shows $\delta E_{g,t}$ (blue), $\delta F_{g,t}$ (violet) and $\delta G_{g,t}$ (magenta) as a function of the time $t$ on the complete duration (CD). The red boxes on the figures represents the 20 min on which the average growth strain is maximal (SD).

The second panel displays the two estimated linear models for the angular distribution of the growth strain rate: the solid line corresponds to the linear model of Equation (3), and the dotted line corresponds to the linear model of Equation (1).

The third panel presents $\delta E_{e,t}$ (red), $\delta F_{e,t}$ (orange) and $\delta G_{e,t}$ (yellow) as functions of the pressure $P$.

The fourth panel shows the two estimated linear models for the angular distribution of the elastic compliance: the solid line corresponds to the linear model of Equation (3), and the dotted line corresponds to the linear model of Equation (1).

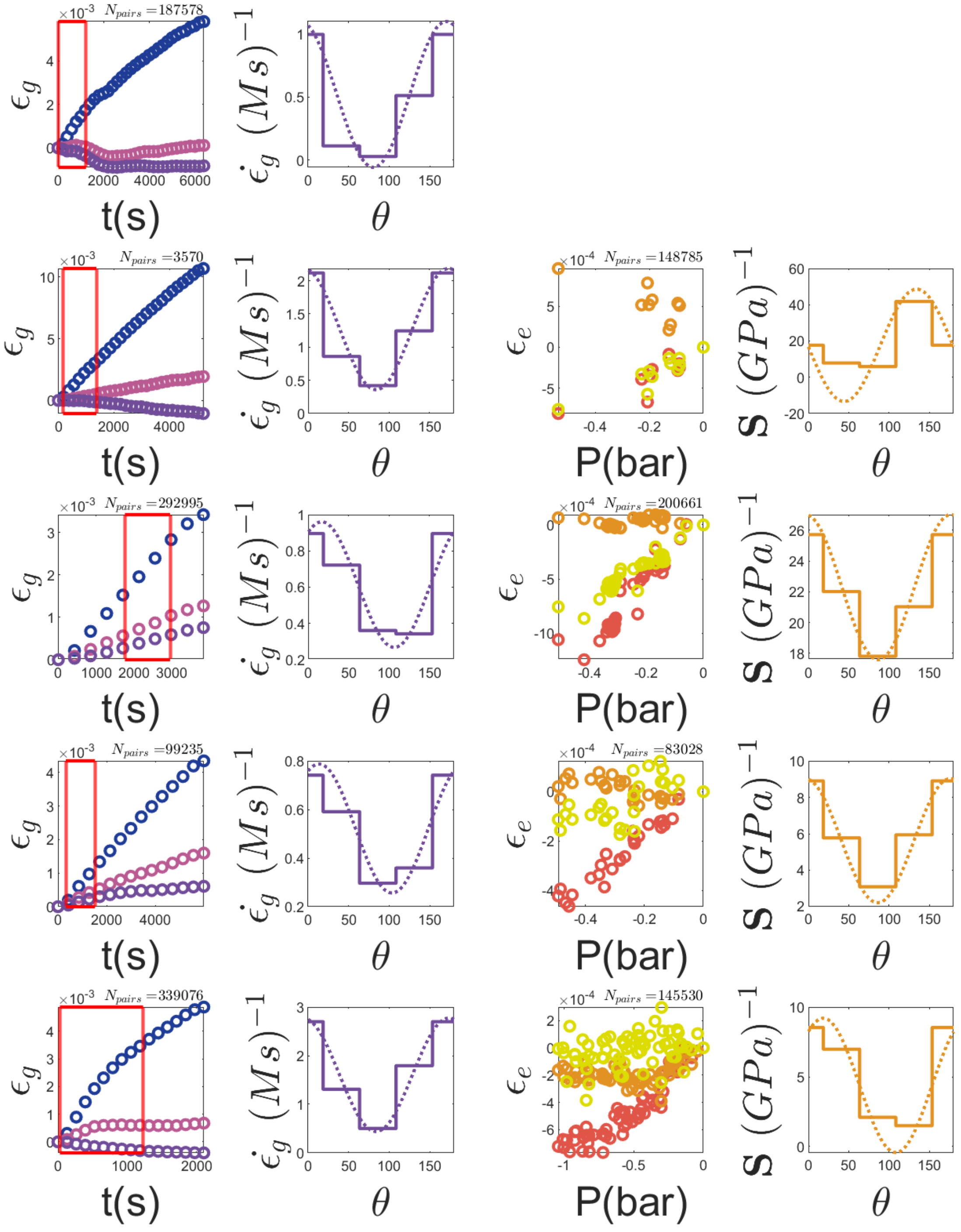
$N_{pairs}$ =187578
$N_{pairs}$ =3570
$N_{pairs}$ =148785
$N_{pairs}$ =292995
$N_{pairs}$ =200661
$N_{pairs}$ =99235
$N_{pairs}$ =83028
$N_{pairs}$ =339076
$N_{pairs}$ =145530
$\epsilon_g$
$\dot{\epsilon}_g\ (Ms)^{-1}$
$\epsilon_e$
$S\ (GPa)^{-1}$
t(s)
$\theta$
P(bar)

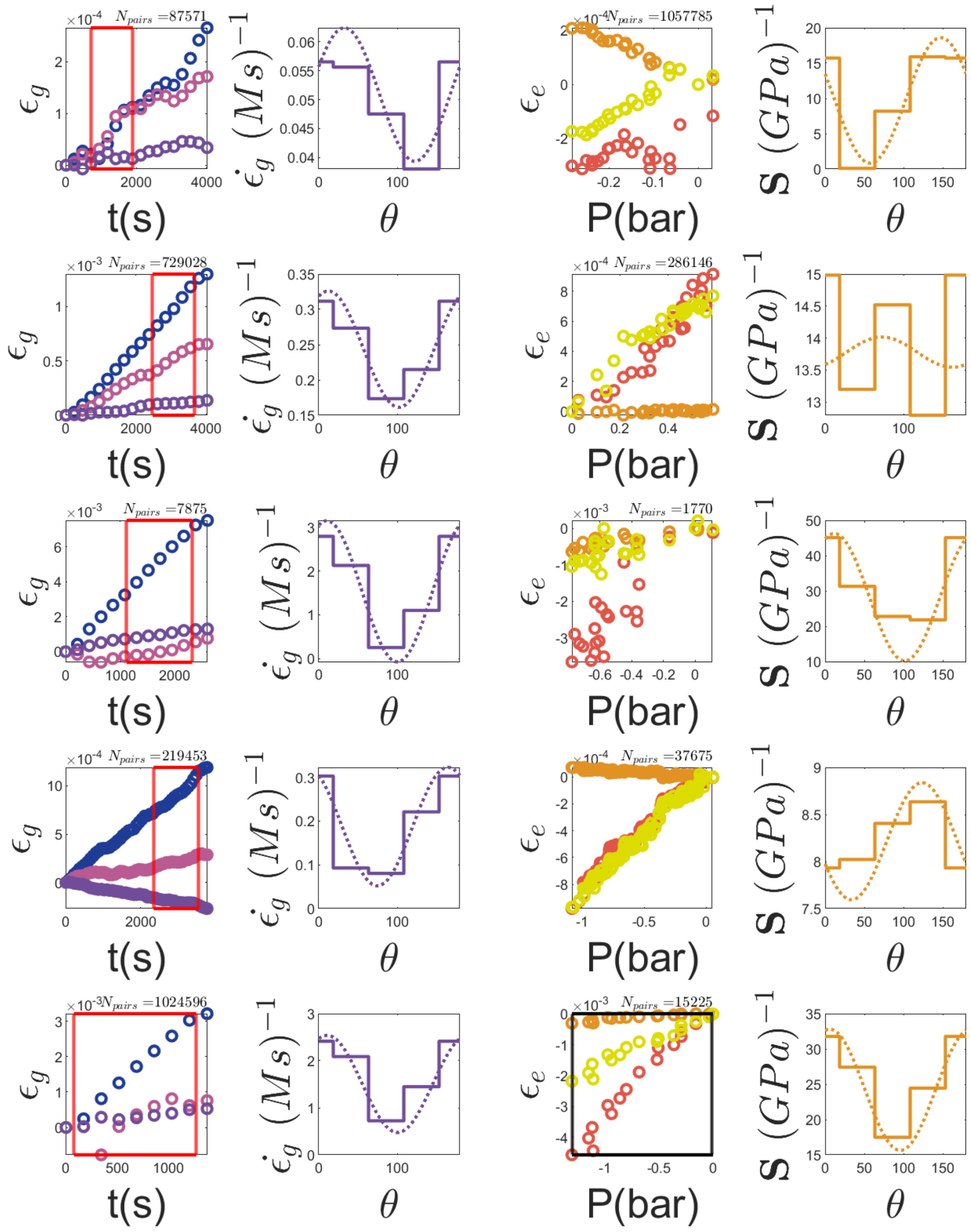

$N_{pairs}$ =87571
$N_{pairs}$ =1057785
$N_{pairs}$ =729028
$N_{pairs}$ =286146
$N_{pairs}$ =7875
$N_{pairs}$ =1770
$N_{pairs}$ =219453
$N_{pairs}$ =37675
$N_{pairs}$ =1024596
$N_{pairs}$ =15225
$\epsilon_g$
$\dot{\epsilon}_g\ (Ms)^{-1}$
$\epsilon_e$
$S\ (GPa)^{-1}$
t(s)
$\theta$
P(bar)

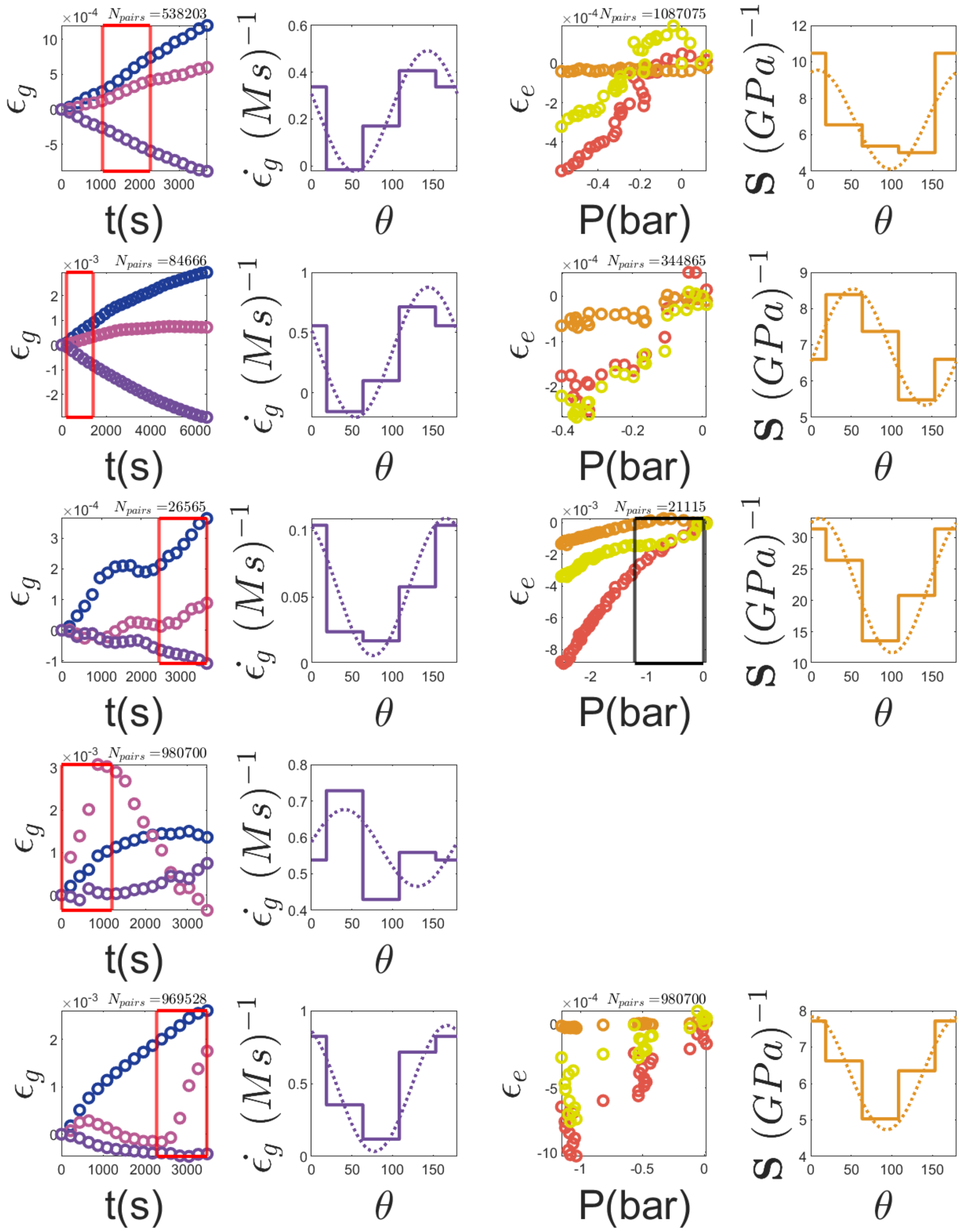

×10-4 Npairs =538203
ϵg
t(s)
ϵ̇g (Ms)−1
θ
×10-4 Npairs =1087075
ϵe
P(bar)
S (GPa)−1
θ
×10-3 Npairs =84666
ϵg
t(s)
ϵ̇g (Ms)−1
θ
×10-4 Npairs =344865
ϵe
P(bar)
S (GPa)−1
θ
×10-4 Npairs =26565
ϵg
t(s)
ϵ̇g (Ms)−1
θ
×10-3 Npairs =21115
ϵe
P(bar)
S (GPa)−1
θ
×10-3 Npairs =980700
ϵg
t(s)
ϵ̇g (Ms)−1
θ
×10-3 Npairs =969528
ϵg
t(s)
ϵ̇g (Ms)−1
θ
×10-4 Npairs =980700
ϵe
P(bar)
S (GPa)−1
θ

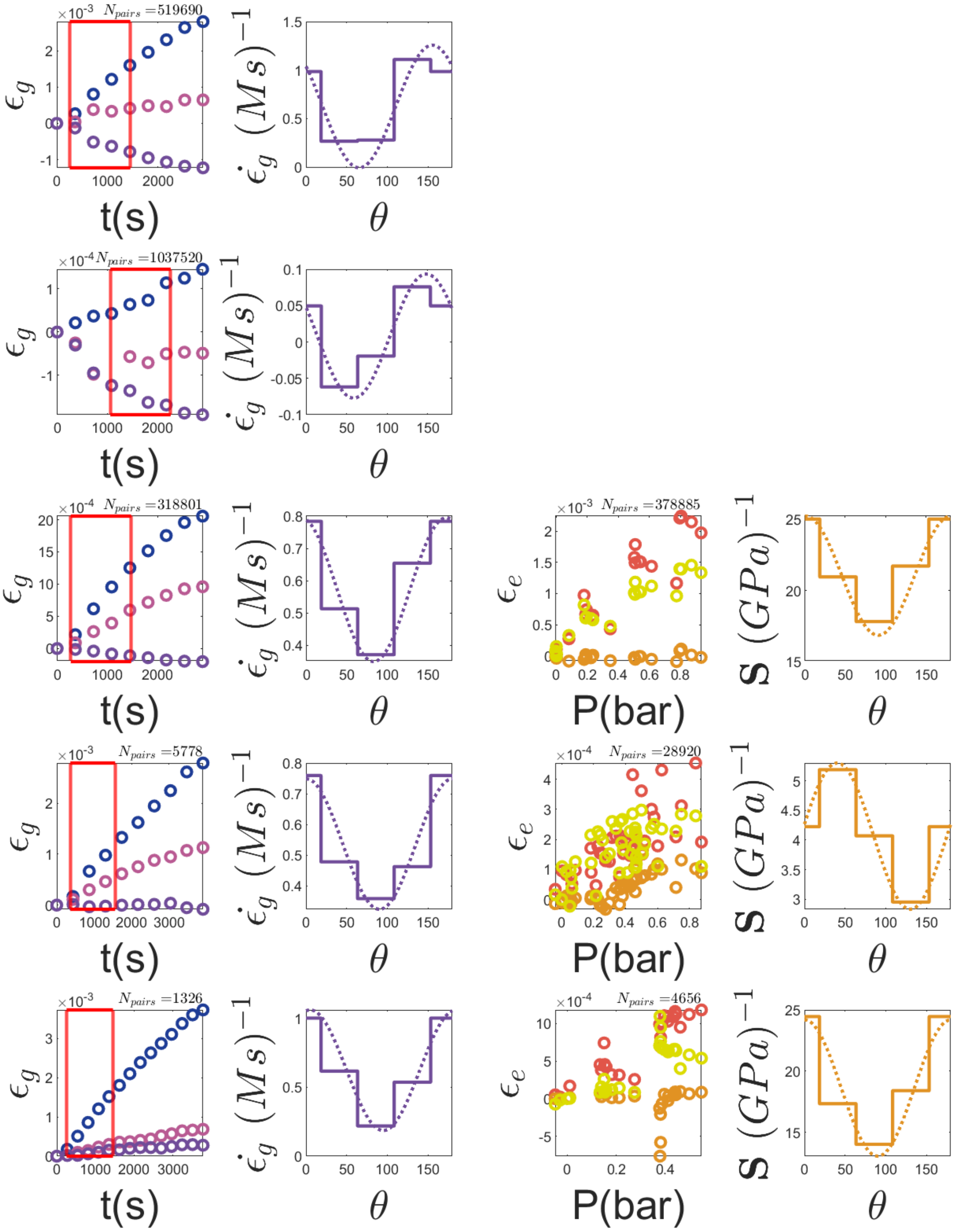

$\times 10^{-3}$ $N_{pairs}$ =519690
$\times 10^{-4}$ $N_{pairs}$ =1037520
$\times 10^{-4}$ $N_{pairs}$ =318801
$\times 10^{-3}$ $N_{pairs}$ =378885
$\times 10^{-3}$ $N_{pairs}$ =5778
$\times 10^{-4}$ $N_{pairs}$ =28920
$\times 10^{-3}$ $N_{pairs}$ =1326
$\times 10^{-4}$ $N_{pairs}$ =4656
$\epsilon_g$
$\epsilon_e$
$\dot{\epsilon}_g\ (M\,s)^{-1}$
$\mathbf{S}\ (GPa)^{-1}$
t(s)
$\theta$
P(bar)

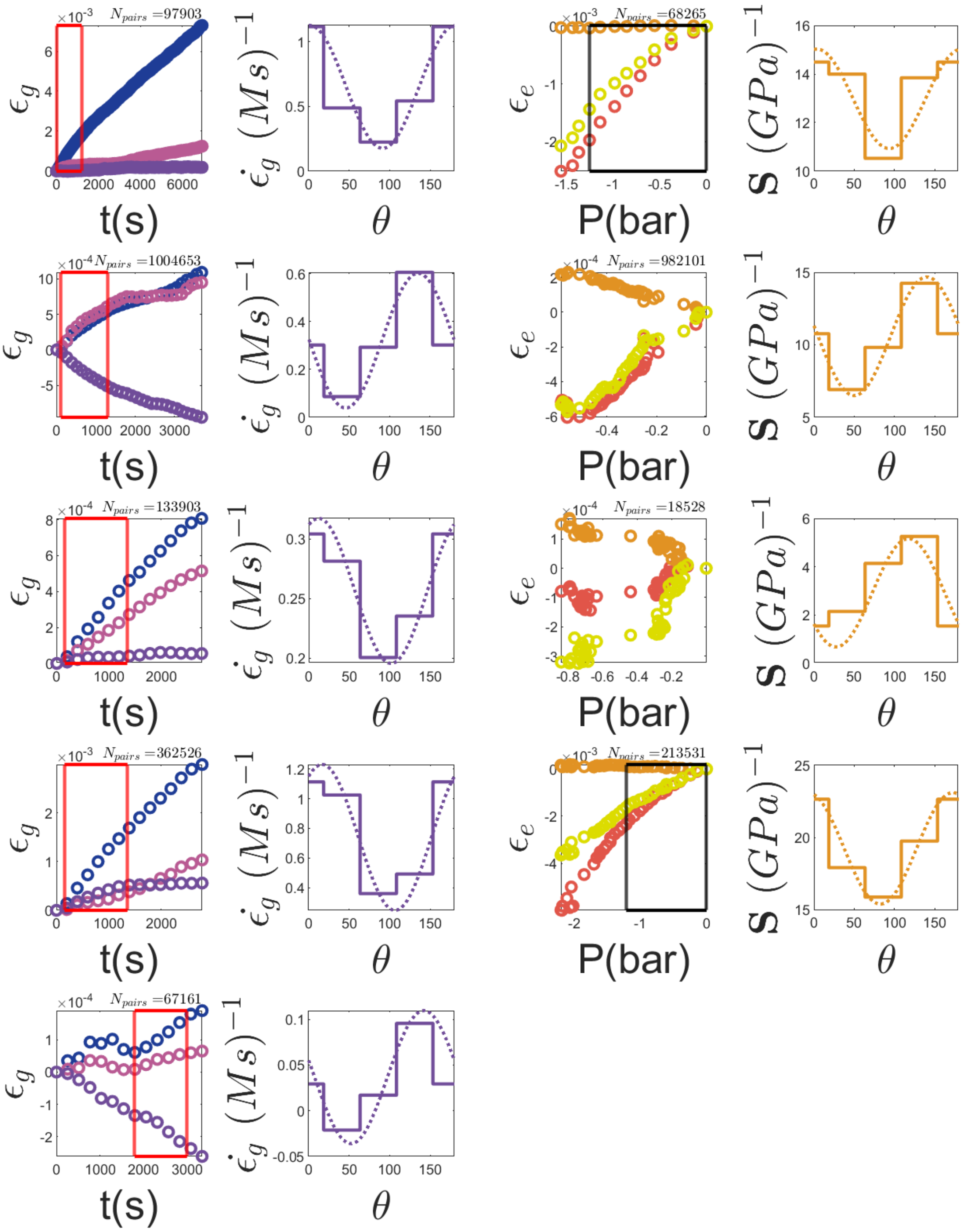

$N_{pairs}$ =97903
$N_{pairs}$ =68265
$N_{pairs}$ =1004653
$N_{pairs}$ =982101
$N_{pairs}$ =133903
$N_{pairs}$ =18528
$N_{pairs}$ =362526
$N_{pairs}$ =213531
$N_{pairs}$ =67161
$\epsilon_g$
$\epsilon_e$
$\dot{\epsilon}_g\ (Ms)^{-1}$
$S\ (GPa)^{-1}$
t(s)
$\theta$
P(bar)

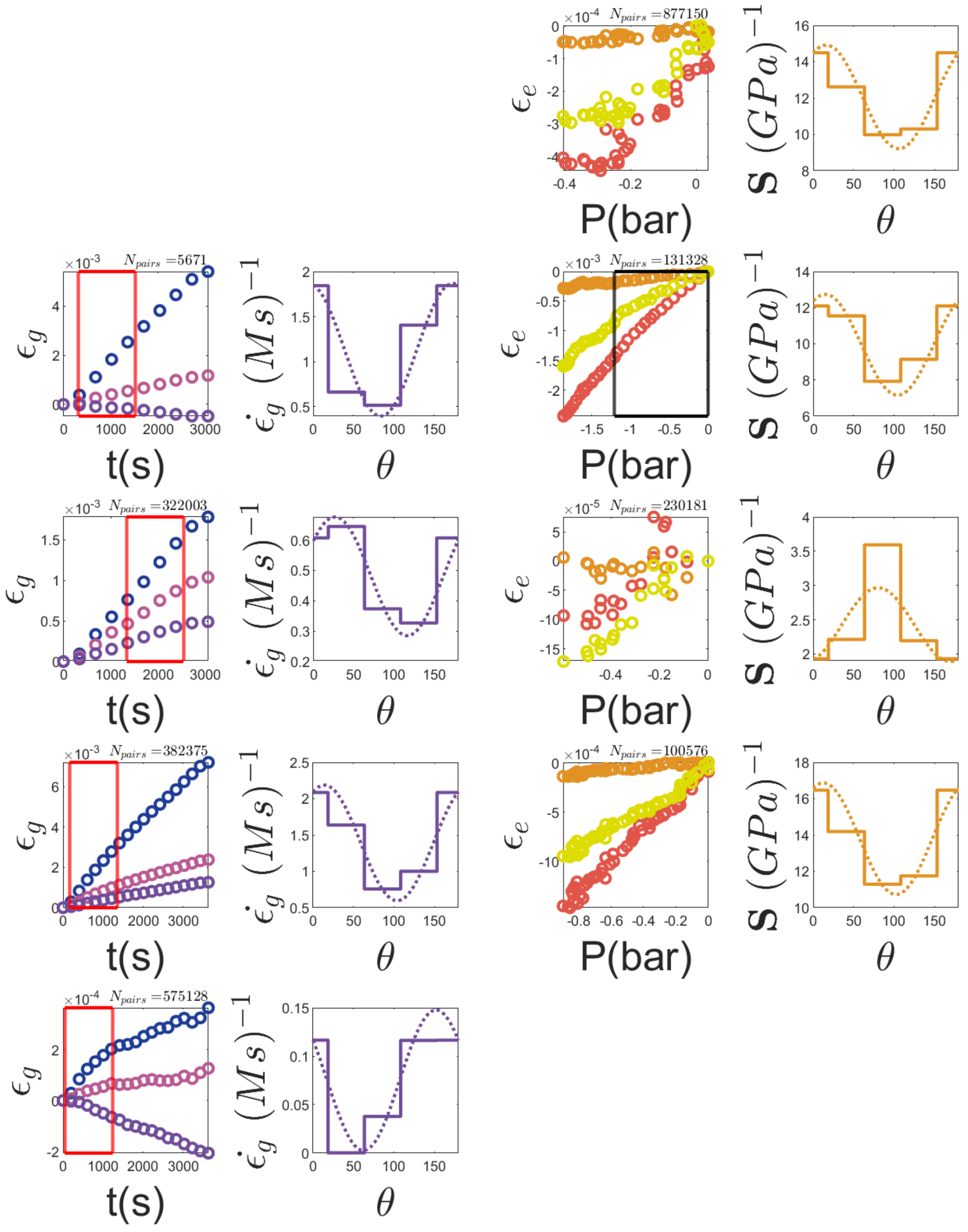

$N_{pairs}$ =877150
$N_{pairs}$ =5671
$N_{pairs}$ =131328
$N_{pairs}$ =322003
$N_{pairs}$ =230181
$N_{pairs}$ =382375
$N_{pairs}$ =100576
$N_{pairs}$ =575128
$\epsilon_g$
$\epsilon_e$
t(s)
P(bar)
$\theta$
$\dot{\epsilon}_g\ (Ms)^{-1}$
$S\ (GPa)^{-1}$

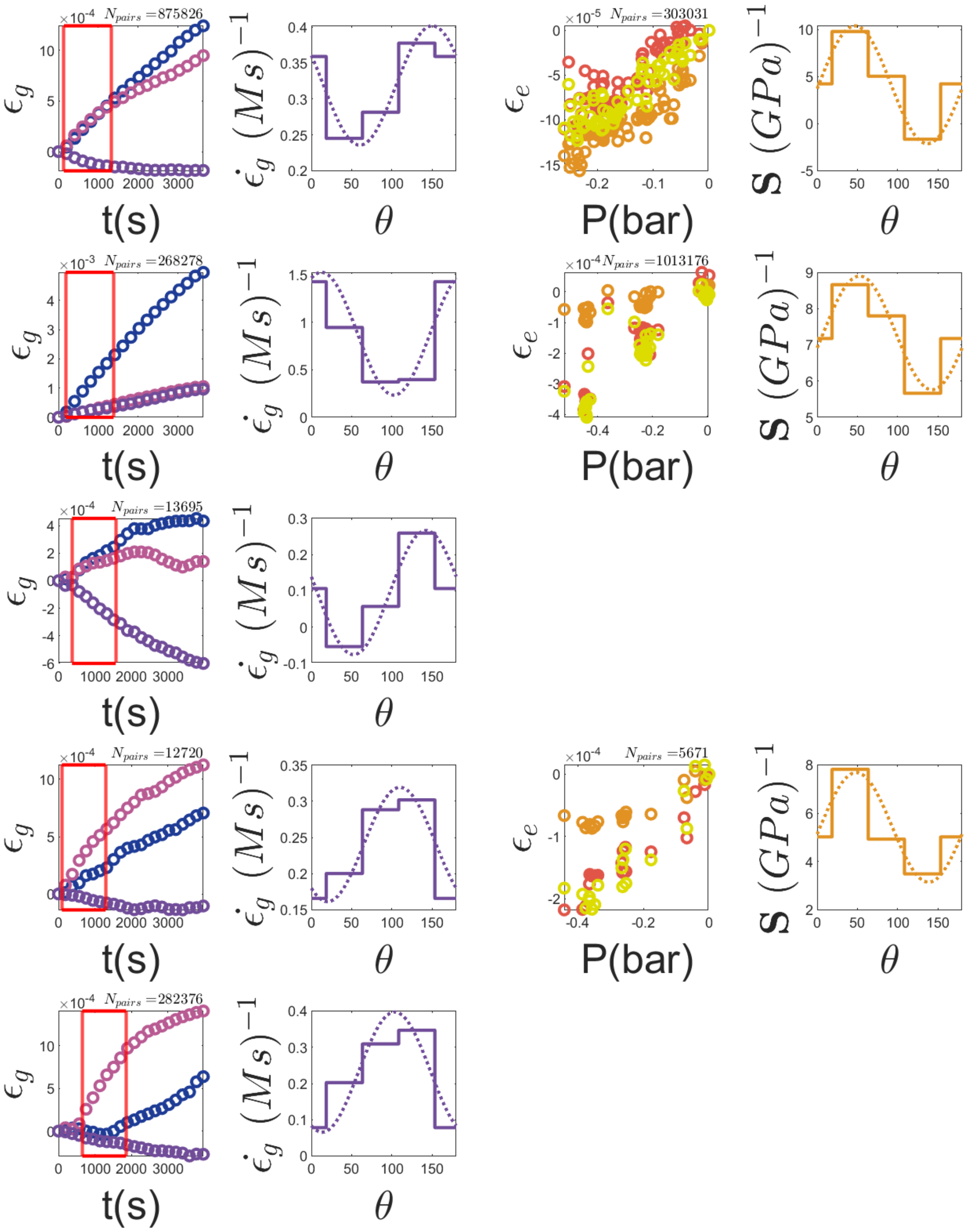
$N_{pairs}=875826$
$N_{pairs}=303031$
$N_{pairs}=268278$
$N_{pairs}=1013176$
$N_{pairs}=13695$
$N_{pairs}=12720$
$N_{pairs}=5671$
$N_{pairs}=282376$
$\epsilon_g$
$\epsilon_e$
t(s)
P(bar)
$\theta$
$\dot{\epsilon}_g\ (Ms)^{-1}$
S $(GPa)^{-1}$

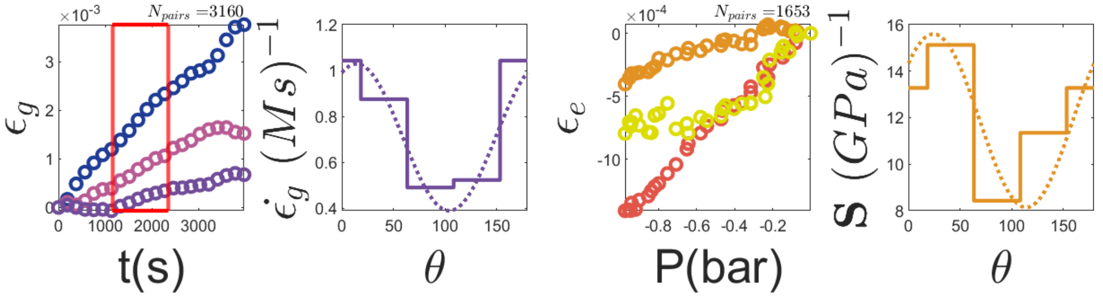


### 4 Correlation tests

#### 4.1 Correlation coefficient obtained for complete duration (CD)

Table 1: Cylinder coordinate system. Spearman $\rho$ correlation coefficients ($N = 28$, elastic compliances; $N = 35$, growth strain rates; $N = 24$, correlations between elastic compliances and growth strain rates). (CD)

| $\rho$ | $R$ | $\mathbf{S}_{\text{ax}}$ | $\mathbf{S}_{45}$ | $\mathbf{S}_{\text{rad}}$ | $\mathbf{S}_{135}$ | $\dot{\epsilon}_{g,\text{ax}}$ | $\dot{\epsilon}_{g,45}$ | $\dot{\epsilon}_{g,\text{rad}}$ | $\dot{\epsilon}_{g,135}$ |
|---|---|---|---|---|---|---|---|---|---|
| $R$ | | $-0.32$ | $-0.30$ | $-0.02$ | $-0.23$ | $-0.65$ | $-0.71$ | $-0.57$ | $-0.45$ |
| $\mathbf{S}_{\text{ax}}$ | | | 0.77 | 0.85 | 0.92 | 0.53 | 0.53 | 0.23 | 0.38 |
| $\mathbf{S}_{45}$ | | | | 0.86 | 0.66 | 0.48 | 0.51 | 0.19 | 0.25 |
| $\mathbf{S}_{\text{rad}}$ | | | | | 0.83 | 0.27 | 0.32 | 0.03 | 0.15 |
| $\mathbf{S}_{135}$ | | | | | | 0.42 | 0.41 | 0.16 | 0.35 |
| $\dot{\epsilon}_{g,\text{ax}}$ | | | | | | | 0.85 | 0.67 | 0.82 |
| $\dot{\epsilon}_{g,45}$ | | | | | | | | 0.83 | 0.60 |
| $\dot{\epsilon}_{g,\text{rad}}$ | | | | | | | | | 0.63 |

Table 2: Cylinder coordinate system. $p$-values of Spearman correlation coefficients ($N = 28$, elastic compliances; $N = 35$, growth strain rates; $N = 24$ correlations between elastic compliances and growth strain rates). (CD)

| $p$-value | $R$ | $\mathbf{S}_{\rm ax}$ | $\mathbf{S}_{45}$ | $\mathbf{S}_{\rm rad}$ | $\mathbf{S}_{135}$ | $\dot\epsilon_{g,\rm ax}$ | $\dot\epsilon_{g,45}$ | $\dot\epsilon_{g,\rm rad}$ | $\dot\epsilon_{g,135}$ |
|---|---|---|---|---|---|---|---|---|---|
| $R$ | | 0.10 | 0.12 | 0.93 | 0.23 | $3\times10^{-5}$ | $5\times10^{-6}$ | 0.0005 | 0.007 |
| $\mathbf{S}_{\rm ax}$ | | | $4\times10^{-6}$ | $10^{-6}$ | $5\times10^{-7}$ | 0.009 | 0.008 | 0.28 | 0.06 |
| $\mathbf{S}_{45}$ | | | | $10^{-6}$ | 0.0002 | 0.02 | 0.01 | 0.36 | 0.23 |
| $\mathbf{S}_{\rm rad}$ | | | | | $10^{-6}$ | 0.19 | 0.13 | 0.87 | 0.48 |
| $\mathbf{S}_{135}$ | | | | | | 0.04 | 0.05 | 0.45 | 0.10 |
| $\dot\epsilon_{g,\rm ax}$ | | | | | | | $9\times10^{-8}$ | $2\times10^{-5}$ | $2\times10^{-7}$ |
| $\dot\epsilon_{g,45}$ | | | | | | | | $2\times10^{-7}$ | 0.0002 |
| $\dot\epsilon_{g,\rm rad}$ | | | | | | | | | $7\times10^{-5}$ |

Table 3: Principal direction. Spearman $\rho$ correlation coefficient ($N = 28$, elastic compliances; $N = 35$, growth strain rates; $N = 24$ correlations between elastic compliances and growth strain rates). (CD)

| $\rho$ | $R$ | $\theta_{\perp,e}$ | $\mathbf{S}_\perp$ | $\mathbf{S}_\parallel$ | $\theta_{\perp,g}$ | $\dot\epsilon_{g,\perp}$ | $\dot\epsilon_{g,\parallel}$ |
|---|---|---|---|---|---|---|---|
| $R$ | | −0.16 | −0.11 | −0.19 | −0.34 | −0.50 | −0.58 |
| $\theta_{\perp,e}$ | | | −0.01 | 0.02 | −0.11 | 0.38 | 0.44 |
| $\mathbf{S}_\perp$ | | | | 0.42 | 0.17 | 0.10 | 0.26 |
| $\mathbf{S}_\parallel$ | | | | | 0.18 | 0.06 | 0.45 |
| $\theta_{\perp,g}$ | | | | | | 0.63 | 0.17 |
| $\dot\epsilon_{g,\perp}$ | | | | | | | 0.34 |

Table 4: Principal direction. Spearman $p$-values ($N = 28$, elastic compliances; $N = 35$, growth strain rates; $N = 24$ correlations between elastic compliances and growth strain rates). (CD)

| $p - value$ | $R$ | $\theta_{\perp,e}$ | $\mathbf{S}_\perp$ | $\mathbf{S}_\parallel$ | $\theta_{\perp,g}$ | $\dot\epsilon_{g,\perp}$ | $\dot\epsilon_{g,\parallel}$ |
|---|---|---|---|---|---|---|---|
| $R$ | | 0.43 | 0.59 | 0.32 | 0.04 | 0.002 | 0.0004 |
| $\theta_{\perp,e}$ | | | 0.95 | 0.91 | 0.62 | 0.07 | 0.03 |
| $\mathbf{S}_\perp$ | | | | 0.03 | 0.42 | 0.65 | 0.23 |
| $\mathbf{S}_\parallel$ | | | | | 0.40 | 0.78 | 0.03 |
| $\theta_{\perp,g}$ | | | | | | $8\times10^{-5}$ | 0.33 |
| $\dot\epsilon_{g,\perp}$ | | | | | | | 0.04 |

#### 4.2 Correlation coefficient obtained for shorter interval (SD)

Table 5: Cylinder coordinate. Spearman $\rho$ correlation coefficients ($N = 28$, elastic compliances; $N = 35$, growth strain rates; $N = 24$, correlations between elastic compliances and growth strain rates). (SD)

| $\rho$ | $R$ | $\mathbf{S}_{\text{ax}}$ | $\mathbf{S}_{45}$ | $\mathbf{S}_{\text{rad}}$ | $\mathbf{S}_{135}$ | $\dot{\epsilon}_{g,\text{ax}}$ | $\dot{\epsilon}_{g,45}$ | $\dot{\epsilon}_{g,\text{rad}}$ | $\dot{\epsilon}_{g,135}$ |
|---|---|---|---|---|---|---|---|---|---|
| $R$ | | −0.32 | −0.30 | −0.02 | −0.23 | −0.51 | −0.64 | −0.41 | −0.30 |
| $\mathbf{S}_{ax}$ | | | 0.77 | 0.85 | 0.92 | 0.37 | 0.43 | 0.18 | 0.27 |
| $\mathbf{S}_{45}$ | | | | 0.86 | 0.66 | 0.32 | 0.37 | 0.13 | 0.21 |
| $\mathbf{S}_{rad}$ | | | | | 0.83 | 0.14 | 0.18 | 0.02 | 0.09 |
| $\mathbf{S}_{135}$ | | | | | | 0.31 | 0.35 | 0.17 | 0.23 |
| $\dot{\epsilon}_{g,ax}$ | | | | | | | 0.85 | 0.80 | 0.88 |
| $\dot{\epsilon}_{g,45}$ | | | | | | | | 0.80 | 0.67 |
| $\dot{\epsilon}_{g,rad}$ | | | | | | | | | 0.89 |

Table 6: Cylinder coordinate. Spearman $p$-values ($N = 28$, elastic compliances; $N = 35$, growth strain rates; $N = 24$, correlations between elastic compliances and growth strain rates). (SD)

| $p$-value | $R$ | $\mathbf{S}_{\text{ax}}$ | $\mathbf{S}_{45}$ | $\mathbf{S}_{\text{rad}}$ | $\mathbf{S}_{135}$ | $\dot{\epsilon}_{g,\text{ax}}$ | $\dot{\epsilon}_{g,45}$ | $\dot{\epsilon}_{g,\text{rad}}$ | $\dot{\epsilon}_{g,135}$ |
|---|---|---|---|---|---|---|---|---|---|
| $R$ | | 0.10 | 0.12 | 0.93 | 0.23 | 0.002 | $5 \times 10^{-5}$ | 0.02 | 0.08 |
| $\mathbf{S}_{ax}$ | | | $4 \times 10^{-6}$ | $\times 10^{-6}$ | $5 \times 10^{-7}$ | 0.07 | 0.03 | 0.40 | 0.20 |
| $\mathbf{S}_{45}$ | | | | $1 \times 10^{-6}$ | 0.0002 | 0.13 | 0.07 | 0.54 | 0.33 |
| $\mathbf{S}_{rad}$ | | | | | $1 \times 10^{-6}$ | 0.50 | 0.39 | 0.93 | 0.67 |
| $\mathbf{S}_{135}$ | | | | | | 0.14 | 0.09 | 0.42 | 0.27 |
| $\dot{\epsilon}_{g,ax}$ | | | | | | | $6 \times 10^{-8}$ | $3 \times 10^{-7}$ | $3 \times 10^{-9}$ |
| $\dot{\epsilon}_{g,45}$ | | | | | | | | $3 \times 10^{-7}$ | $2 \times 10^{-5}$ |
| $\dot{\epsilon}_{g,rad}$ | | | | | | | | | 0 |

Table 7: Principal direction. Spearman $\rho$ ($N = 28$, elastic compliances; $N = 35$, growth strain rates; $N = 24$, correlations between elastic compliances and growth strain rates). (SD)

| $\rho$ | $R$ | $\theta_{\perp,e}$ | $\mathbf{S}_\perp$ | $\mathbf{S}_\parallel$ | $\theta_{\perp,g}$ | $\dot{\epsilon}_{g,\perp}$ | $\dot{\epsilon}_{g,\parallel}$ |
|---|---|---|---|---|---|---|---|
| $R$ | | $-0.16$ | $-0.11$ | $-0.19$ | $-0.41$ | $-0.45$ | $-0.51$ |
| $\theta_{\perp,e}$ | | | $-0.01$ | 0.02 | $-0.23$ | 0.14 | 0.33 |
| $\mathbf{S}_\perp$ | | | | 0.42 | $-0.007$ | $-0.05$ | 0.23 |
| $\mathbf{S}_\parallel$ | | | | | 0.29 | 0.10 | 0.21 |
| $\theta_{\perp,g}$ | | | | | | 0.58 | 0.18 |
| $\dot{\epsilon}_{g,\perp}$ | | | | | | | 0.44 |

Table 8: Principal direction. $P$-values of Spearman correlation coefficients ($N = 28$, elastic compliances; $N = 35$, growth strain rates; $N = 24$, correlations between elastic compliances and growth strain rates)). (SD)

| $p$-value | $R$ | $\theta_{\perp,e}$ | $\mathbf{S}_\perp$ | $\mathbf{S}_\parallel$ | $\theta_{\perp,g}$ | $\dot{\epsilon}_{g,\perp}$ | $\dot{\epsilon}_{g,\parallel}$ |
|---|---|---|---|---|---|---|---|
| $R$ | | 0.43 | 0.59 | 0.32 | 0.02 | 0.007 | 0.002 |
| $\theta_{\perp,e}$ | | | 0.95 | 0.91 | 0.27 | 0.53 | 0.12 |
| $\mathbf{S}_\perp$ | | | | 0.03 | 0.98 | 0.82 | 0.29 |
| $\mathbf{S}_\parallel$ | | | | | 0.16 | 0.64 | 0.33 |
| $\theta_{\perp,g}$ | | | | | | 0.0003 | 0.30 |
| $\dot{\epsilon}_{g,\perp}$ | | | | | | | 0.009 |

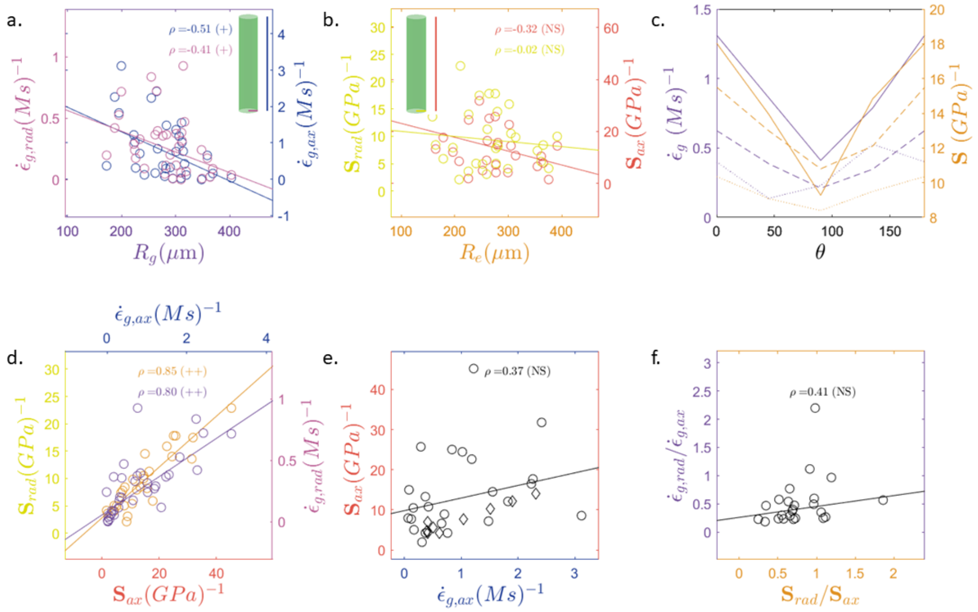

Figure 3: **a**. Radial growth strain rate (magenta) and axial growth strain (blue) vs. cell radius $R_g$ ($N = 35$). **b**. Radial elastic compliance (yellow) and axial elastic compliance (red) vs. cell radius $R_e$ ($N = 28$). **c**. Data pooled into three quantiles based on cell radius ($N = 28$, elastic compliance; $N = 35$, growth strain rate). The first quantile is shown as a solid line, the second as a dashed line, and the third as a dotted line. Right axis (orange): average angular distribution of elastic compliance for each quantile. Left axis (violet): average angular distribution of growth strain rate for each quantile. **d**. Left axis (orange): axial elastic compliance vs. radial elastic compliance ($N = 28$). Right axis (violet): axial growth strain rate vs. radial growth strain rate ($N = 35$). **e**. Axial elastic compliance vs. axial growth strain rate ($N = 24$). (o) Our Data. ($\diamond$) Data extracted from [14]: The growth velocity reported in this reference was converted into a growth strain rate by assuming an internode length of 1 cm, as stated elsewhere in the same reference. $\rho$: Spearman correlation coefficient. **f**. Ratio of elastic compliances versus ratio of growth strains ($N = 24$). In all panels, regression lines were computed using MATLAB's `fitlm` function with the robust `'bisquare'` option. NS: $p$-value $\geq 0.05$; +: $p$-value $< 0.05$; ++: $p$-value $< 0.001$. The growth strains correspond to the (SD) condition.

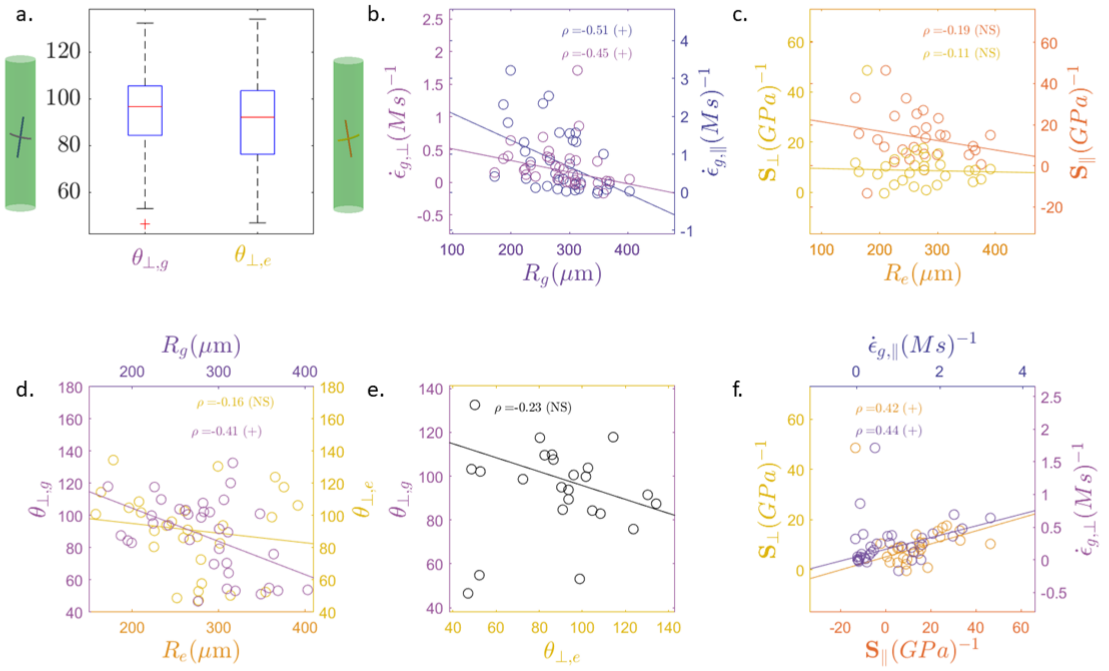


Figure 4: **a**. Boxplots of the principal axis direction of the elastic compliance tensor $\perp$ component (light orange) and the growth strain rate tensor $\perp$ component (light violet)($N = 24$). **b**. Principal growth strain rate in the $\perp$ direction (light violet) and in the $\parallel$ direction (dark violet) vs. cell radius $R_g$ ($N = 35$). **c**. Principal elastic compliance (yellow) in the $\perp$ direction (light orange) and in the $\parallel$ direction (dark orange) vs. cell radius $R_e$ ($N = 28$). **d**. Principal growth strain rate direction (respectively, elastic compliance) in the $\perp$ direction (light violet, respectively light orange) and in the $\parallel$ direction (dark violet, respectively dark orange) vs. cell radius $R_g$ (resp. $R_e$) ($N = 35$, resp. $N = 28$). **e**. Principal growth strain rate in the $\perp$ direction (light violet) vs. principal elastic compliance in the $\perp$ direction (light orange) ($N = 24$). **f**. (Left axis, orange) Elastic compliance in the $\parallel$ direction vs. elastic compliance in the $\perp$ direction ($N = 28$). (Right axis, violet) Growth strain rate in the $\parallel$ direction vs. growth strain rate in the $\perp$ direction ($N = 35$). In all panels, the regression line were computed using MATLAB's `fitlm` function with the option `'robust'` and the `'bisquare'` algorithm. $\rho$ is the Spearman correlation coefficient. NS: $p$-value $\geq 0.05$; +: $p$-value $< 0.05$; ++: $p$-value $< 0.001$. The growth strains correspond to the (SD) condition.
